# The soluble state of the HIV-1 Vpu protein forms a complex with Ca^2+^-calmodulin

**DOI:** 10.1101/2025.06.12.658902

**Authors:** Olamide Ishola, Md Majharul Islam, Elaheh Hadadianpour, Peter P. Borbat, Adeyemi Ogunbowale, Juan C. R. Amador, Elka R. Georgieva

**Author notes:** Corresponding author: Elka R. Georgieva. Equal contribution (Olamide Ishola, Md Majharul Islam).

## Abstract

The HIV-1 Vpu membrane protein is crucial to the virus lifecycle. Our recent studies revealed soluble Vpu oligomers, prompting further investigation into their interactions with cellular proteins. Notably, Vpu may form a complex with calmodulin (CaM) due to its putative CaM-binding motif; however, experimental proof of this association remained unavailable.

Here, we present definitive experimental evidence that the soluble Vpu complex interacts *in vitro* with calcium-bound CaM (Ca^2+^-CaM) its active form. Using double electron electron-resonance (DEER) spectroscopy and protein spin labeling, we detected the formation of a soluble Vpu–Ca^2+^-CaM complex. Both the full-length (FL) and truncated C-terminal regions of Vpu bind Ca^2+^-CaM. DEER experiments on a spin-labeled CaM cysteine mutant S39C/A103C revealed that, upon association with Vpu, Ca^2+^-CaM undergoes a transition from an open to a more closed conformation, consistent with previous reports of Ca^2+^-CaM interactions with other proteins.

Furthermore, we observed that the binding of Vpu to Ca^2+^-CaM leads to dissociation of soluble Vpu oligomers, as evidenced by a reduction in DEER modulation depth for FL Vpu spin-labeled at residue L42C. FRET analysis with a fluorescently labeled C-terminal cysteine mutant of Vpu confirmed this result. Like FL Vpu, the Vpu C-terminal region forms soluble homooligomers that dissociate upon binding to Ca^2+^-CaM. Collectively, our results suggest that soluble Vpu and Ca^2+^-CaM form an equimolar complex. DEER analysis of Vpu C-terminal region spin-labeled at residues Q36C/I61C demonstrated that Vpu undergoes significant conformational changes to facilitate Ca^2+^-CaM binding. These findings could be relevant to Vpu-CaM interactions under physiological conditions.

## INTRODUCTION

Calmodulin (CaM) is a calcium-sensing protein playing key roles in cellular processes, e.g., it regulates the activity of enzymes and ion channels, affects the processes of development and cytoskeletal organization, etc.^1^ In higher organisms, CaM is conserved, being ubiquitously expressed at high concentrations (>0.1% of total protein in the cells) and distributed to all cellular compartments^1, 2^. Extensive studies have identified and characterized several mechanisms of CaM interactions with its protein targets, which usually bind to CaM through electrostatic and hydrophobic interactions alike, both of which are presented by positively-charged and hydrophobic residues in amphipathic helices^3^. Examples of target protein motifs that interact with Ca2+-bound CaM include: 1-10 [FILVW]xxxxxxxx[FILVW], 1-5-10 [FILVW]xxx[FAILVW]xxxx[FILVW], and the basic motif 1-5-10 [RK][RK][RK][FAILVW]xxx[FILV]xxxx[FILVW]. (Herein, x is any aa). Note that CaM interacts with some proteins in Ca^2+^-independent manner. Such proteins contain the following motifs: IQ [FILV]Qxxx[RK]Gxxx[RK]xx[FILVWY], IQ-like [FILV]Qxxx[RK]xxxxxxxx that may require Ca^2+^ in special cases, IQ-2A [IVL]QxxxRxxxx[VL][KR]xW, etc.^4, 5^

Because of the prominent role of CaM in regulating cellular processes through protein-protein interactions, prior studies focusing on HIV-1 have identified and characterized a group of HIV-1 encoded proteins that recruit CaM to carry out their functions in the infected cells. It was well known that the association of CaM with the HIV-1 encoded Nef, gp160, as well as the matrix (MA) domain of Gag proteins, are all essential for virus replication^6–11^. The HIV-1 Tat protein interaction with Ca^2+^-CaM facilitates the cell apoptosis^12^. It was also proposed that the HIV-1 Gag protein could recruit CaM for trafficking toward the plasma membrane to the site of virus assembly, as the CaM and Gag were found to co-localize in the cytosol^9^. In addition to the already characterized direct interactions between HIV-1 proteins and CaM, CaM-binding sites were predicted for most of eighteen HIV-1-encoded proteins^12^. These findings make a compelling case that HIV-1 has developed efficient mechanisms to exploit the ubiquitous presence of CaM in the cell.

This work thus sought to investigate the interaction of the HIV-1 encoded protein Vpu with CaM. The putative CaM-binding motif (AIVVWSIVII<u>EYRKILRQRKIDRLI</u>) of Vpu, which has no strong homology to the well characterized CaM-binding motifs, was predicted more than 10 years ago^12^. However, no prior experimental data confirming the Vpu-CaM direct association has been reported. Vpu is an insufficiently understood accessory protein of HIV-1, playing significant role in antagonizing cellular proteins, such as the immature CD4 receptor and tetherin by preventing them from being delivered to the plasma membrane (PM) through early ubiquitination and degradation and/or by deactivation of these proteins at the PM. Consequently, Vpu aids virus release^13, 14^. At a closer look, Vpu is a small protein having an N-terminal transmembrane (TM) helix (helix 1) followed by soluble helices 2 and 3. The helix 2 is amphipathic with the potential to associate with the membrane surface via electrostatic interactions, whereas the helix 3, being mostly acidic, resides in the aqueous environment (Figures 1A and B)^15^. It was previously believed that Vpu is an exclusively membrane-resident protein, being found in membranes of endoplasmic reticulum (ER), Golgi and in the PM^15–18^. Recently, however, our lab discovered that Vpu can exist as a soluble homooligomeric complex^19, 20^, whose physiological relevance we seek to understand.

**Figure 1.**
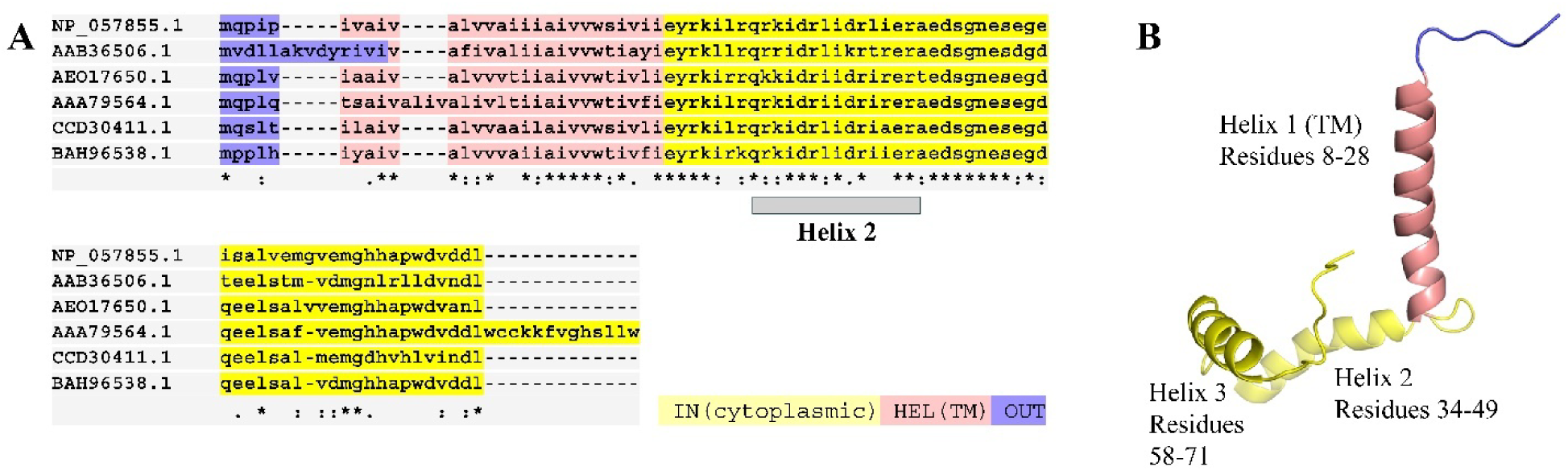
Amino acid (aa) sequence and membrane-bound structure of the HIV-1 Vpu protein: (**A**) The alignment of aa sequences of Vpu protein of randomly selected diverse HIV-1 strains, as follow: NP_057855.1 (strain for reference annotation); AAB36506.1 (an HIV-1 isolate of genetic subtype C); AEO17650.1 (HIV-1 sero-prevalent individual, subtype B); AAA79564.1 (isolate “cntrl 1”, clone=“3”; CCD30411.1 (group M, subgroup G); and BAH96538.1 (multidrug-resistant HIV-1 subtype B). In addition to the high conservation of the TM helix 1 (highlighted in tan), the composition of the loop between helix 1 and helix 2, and the helix 2 and 3 also demonstrates high sequence similarity. The helix 2 is emphasized with gray bar under the aa sequences. (**B**) The solid-state NMR structure of full-length Vpu in lipid (PDB # 2N28) is shown with the helices 1, 2 and 3 indicated and color-coded to correspond to the amino acid sequence shown in (**A**). Helix 1 traverses the membrane bilayer (not shown), whereas helix 2 lays nearly parallel to the membrane surface. The unstructured loops with unknown ordering between helices 1 and 2, and helices 2 and 3 are shown for clarity. The T-COFFEE software was used to align the Vpus’ sequences, and PyMOL was used to visualize the ssNMR structure of Vpu.

To gain insights into the roles of the soluble Vpu, we evaluated its interaction with CaM *in vitro*. We found that it associates with Ca^2+^-CaM and upon this interaction the Ca^2+^-CaM undergoes restructuring to a more closed conformation, similar to previously observed Ca^2+^-CaM complexation with cellular and HIV-1 proteins^4, 12^. Our results support the model which posits that the soluble Vpu-CaM is an equimolar heterocomplex. We have also found that Ca^2+^-CaM interacts with the truncated C-terminal region of Vpu encompassing the residues 29-81 (<u>EYRKILRQRKIDRLI</u>DRLIERAEDSGNESEGEISALVEMGVEMGHHAPWDVDDL) or residues 29-78, which contain only a part (underlined) of the putative CaM-binding site^12^. This may indicate that Vpu has secondary binding sites for CaM, as found for other proteins^5, 21–23^. In this work, we utilized protein engineering to generate spin labeled protein variants needed to obtain distance constraints using pulse dipolar electron spin resonance (ESR) spectroscopy, specifically double electron-electron resonance (DEER). We further applied FRET spectroscopy to fluorescently labeled Vpu C-terminal region and CaM to probe the Vpu soluble oligomer dissociation upon binding to Ca^2+^-CaM. The conformational alterations in Vpu and Ca^2+^-CaM were probed by measuring DEER distances between nitroxide spin label pairs attached to engineered cysteine residues within these proteins. We also conducted amino acid sequence analysis to identify the putative binding motifs in the C-terminal region of Vpu, which could contribute to the interaction with CaM.

A reasonable conjecture based on our results is that under physiological conditions a soluble Vpu-Ca^2+^-CaM complex is formed. Further hypothesis could be that this complex may serve as a vehicle for trafficking Vpu to its membrane destination, which is conceptually analogous to the proposed role of the Gag-CaM association^9^. Our findings contribute to the understanding of HIV-1 Vpu protein interactions with cellular components.

## RESULTS

### Analysis of the putative CaM-binding sites in the Vpu protein—localized in the TM helix 1, amphipathic helix 2, and their interconnecting loop

Because of the previously suggested putative CaM-binding motif in Vpu^12^, we carried out the analysis of Vpu amino acid (aa) sequence with the goal to identify regions which contain the residues found in the motifs of CaM-binding proteins. We focused on the aa composition of the proposed CaM-binding region (Figure 2 A) but also have expanded the search to account for residues in the helix 2 of Vpu (Figure 2 B). As in cases of other CaM-binding proteins encoded by HIV-1^9, 12^, we can identify short sequences of aa residues in Vpu that are identical to some of the canonical CaM-binding motifs^4, 24^. Noteworthy, one can recognize that Vpu sequence has aa regions, which are similar to more than one of these known motifs (Figure 2C). Hence, the earlier proposed CaM-interacting region in Vpu has regions close to the patches of hydrophobic aa’s in the 1-8-14, 1-5-10 and IQ-like CaM-binding motifs. These hydrophobic residues are mostly located in the TM helix 1 of Vpu (Figure 2), i.e. IVVW sequence, highlighted in yellow in Figure 2A. On the other hand, the region of the Vpu helix 2 has more resemblance to the 1-5-10 and IQ-like CaM-binding motifs due to the pairs of positive residues RK, highlighted in magenta in Figure 2A and B. Furthermore, Vpu has a conserved Q (Gln) residue right before the helix 2 (highlighted in cyan in Figure 2A and B), which conceivably acts like the conserved Q residue in the IQ-like motif found in PEP-19 and RC3 proteins^25^.

**Figure 2.**
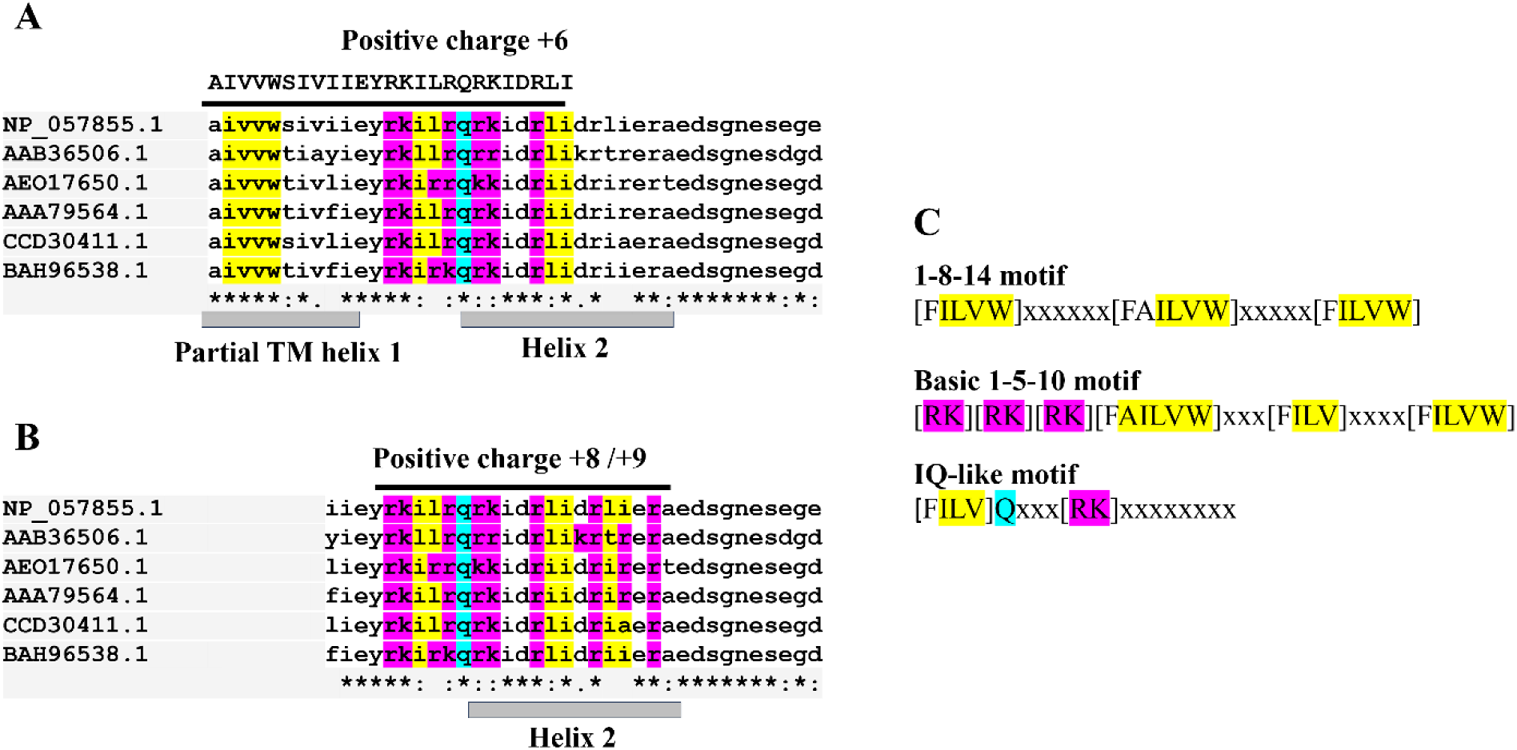
Analysis of the CaM-binding sites in the Vpu aa sequence. (**A**) The alignment of Vpu aa sequences corresponding to the protein region containing the partial TM helix 1, helix 2 and the loops between helices 1, 2 and 2, 3. The aa sequence of the previously predicted CaM-binding motif and its positive charge (+6) are shown above the aligned sequences. (**B**) The aligned Vpu sequences corresponding to the protein region containing the helix 2 and the loops between helices 1, 2 and 2, 3. The positive charge of this aa region (+8 or +9, depending on the HIV-1 strain) is shown. For both (A) and (B) cases the Vpu sequences are from the HIV-1 strains compiled in Figure 1. (**C**) The sequences of the common CaM-binding motifs, which have resemblance with the sequence of the putative Vpu’s CaM-binding region. The “x” in these sequences denotes any aa. Proteins posessing such motifs bind CaM in Ca^2+^-dependent manner, with the exclusion of IQ-like motif which could bind irrespectively of Ca^2+^. In all these aa sequences, the basic residues, as those found in the basic 1-5-10 motif of CaM-binding proteins, are highlighted in magenta. The hydrophobic residues, like those found in the 1-8-14 and/or 1-14 motifs in CaM-binding proteins are highlighted in yellow. The conserved Gln (Q) residue in Vpu aa sequences, which could be like the Gln residue found in the IQ-like motif of CaM-binding proteins is highlighted in cyan. It may well be that Vpu possesses a hybrid motif to associate with CaM in special way via both electrostatic and hydrophobic interactions.

Summarizing the sequence analysis, we note that Vpu possesses a hybrid CaM-binding sequence (motif) or possibly two distinct motifs interacting with CaM via electrostatic and hydrophobic interactions. Furthermore, the basic 1-5-10 and 1-8-14 motifs associate with CaM in a Ca^2+^-dependent manner^4, 24^, which suggested that Vpu could also interacts with Ca^2+^-CaM.

### DEER spectroscopy shows that the Ca^2+^-bound CaM adopts more closed conformation upon binding to the Vpu soluble form

Prior studies have shown that CaM is a binding partner of other HIV-1 encoded proteins^8, 12^, and the HIV-1 protein Vpu has a predicted CaM-binding sequence which resembles the canonical sequences interaction with Ca^2+^-CaM^12^ (Figure 2). Moreover, Vpu can be in a soluble form^19, 20^. Here, we studied the Vpu-Ca^2+^-CaM interaction using highly purified full-length (FL) and truncated C-terminal region of Vpu constructs for binding CaM (Figure S4). As a technicality, both Vpu constructs were fused to a SUMO tag as shown in Figures S1, S2, and S3.

As noted above, our primary goal to characterize the conformational changes taking place in Ca^2+^-CaM upon interaction with the soluble Vpu was addressed by carrying out DEER measurements of Ca^2+^-CaM, doubly spin-labeled with methanethiosulfonate (MTS) nitroxide spin label, MTSL, coupled to the engineered cysteine residues S39C/A103C (Figure 3). The residues reside in the respective globular domains, N- and C-terminal lobes, separated by a long central helix in the Ca^2+^-CaM (Figure 3A). DEER signals for this CaM variant were recorded for Ca^2+^-CaM without Vpu added and in the presence of unlabeled soluble SUMO-FL Vpu and SUMO-Vpu C-terminal region using CaM-to-Vpu variant molar ratio of 1:0.75(0.8). All samples contained 1 mM Ca^2+^. It was expected that free CaM (no Vpu added) was predominantly in its well-defined extended Ca^2+^-bound conformation^2^. This was confirmed by recording the DEER signal trace showing conspicuous oscillations (Figure 3B, upper panel). This well-defined deeply oscillating DEER signal was reconstructed into a narrow distance distribution with a maximum at about 5.3 nm between the labels in good agreement with the expected 4.1 nm distance between the C*_β_* atoms for the residues S38 and A102 in the crystal structure, corresponding to the distance between S39 and A103 in our sequence numbering (Figure 3A, upper panel). Note that our CaM construct was one residue longer than those whose crystal structures were solved and used in this study (PDB # 1EXR and PDB # 1IQ5) to compare the structure-predicted vs. experimental DEER distances. We used the macromolecule modeling software MMM^26^ to model the distance distribution between the MTSL labels attached to S39C and A103C residues, based on generated MTSL rotamer libraries, and found that the model agrees very well with the experimental DEER distances (Figure S5). The longer DEER distance with the maximum at 5.3 nm is well in line with the experimental results and–considering MTSL side-chain size and label orientations–predicted C*_β_*-C*_β_* distance, as noted above. Furthermore, the DEER modulation depth (DEER signal amplitude at zero evolution time in the background-corrected “dipolar” signal) in this measurement corresponded exactly to the calculation for a pair of nitroxide spin labels in a fully labeled protein at the DEER setup used in this work^27–30^. As expected, no distances were reported by DEER for Ca^2+^-CaM with a single spin-label at residue S39C (Figure S6). These results suggest that only well-defined Ca^2+^-CaM monomer in extended conformation was present in the sample.

**Figure 3.**
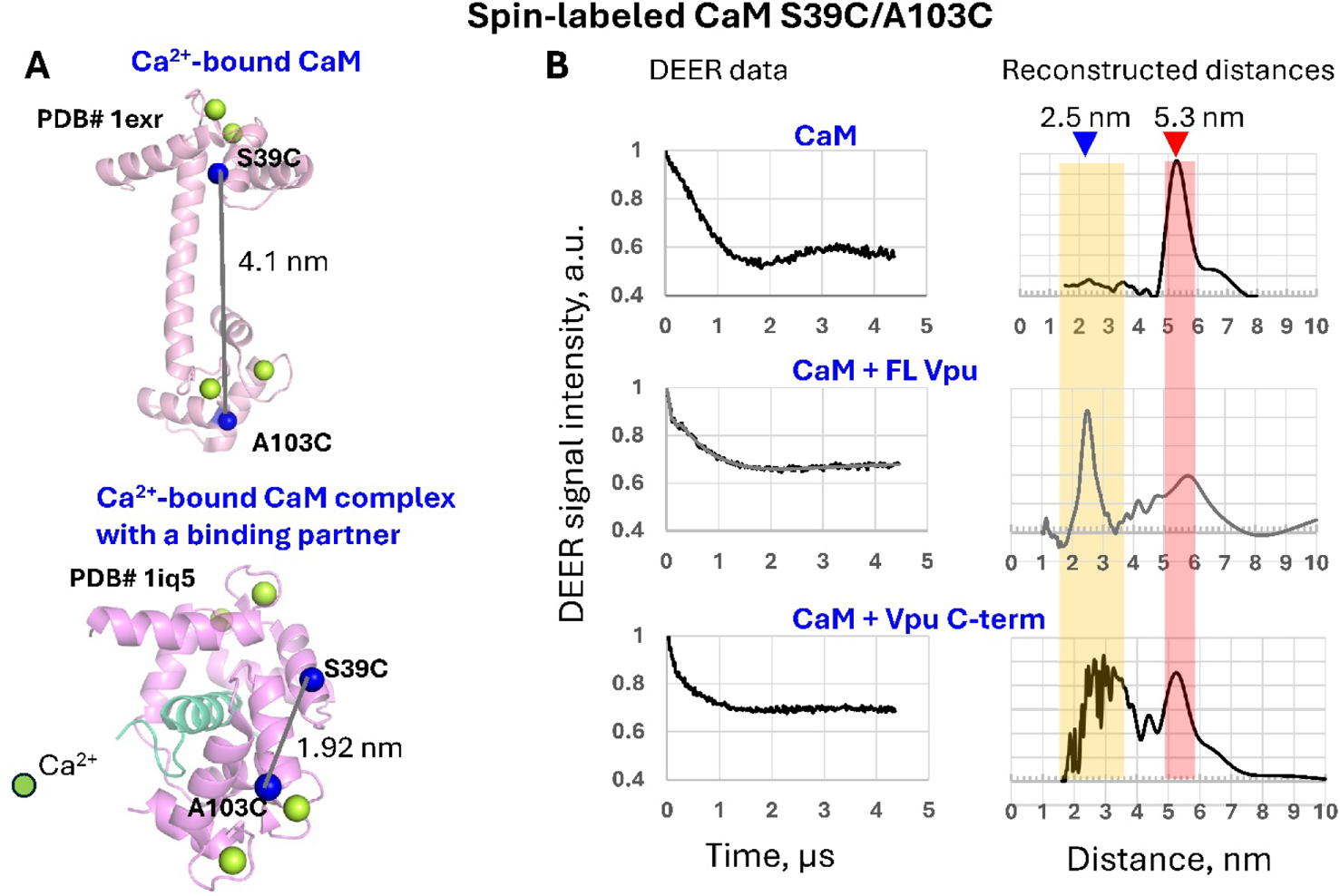
Ca^2+^-CaM interacts with the FL Vpu and truncated C-terminus of Vpu. (**A**) The crystal structures of Ca^2+^-bound CaM (PDB # 1EXR) and Ca^2+^-bound CaM in the complex with a calmodulin dependent kinase fragment (cyan) (PDB # 1IQ5) illustrate extended and closed conformations, respectively. The C*_β_*-atoms of residues S39 and A103 are shown as blue spheres. In the structures shown, these residues are S38 and A102, since these CaM forms are shorter by 1 aa compared to the form we used. The estimated C*_β_*-C*_β_* distances are also shown. (**B**) Baseline-slope corrected DEER data (left), accompanied by the reconstructed inter-spin distances (right) for the following cases: (**B**, **top**) free Ca^2+^-CaM, doubly spin-labeled at residues S39C/A103C; (**B**, **mid**) Ca^2+^-CaM in the presence of an excess of SUMO-FL Vpu; and (**B**, **bottom**) Ca^2+^-CaM in the presence of an excess of the SUMO-Vpu C-terminal region. In the absence of the Vpu variants, distances with a distinct peak at 5.3 nm were obtained between the spin-labeled residues 39 and 103 for Ca^2+^-CaM. However, with the addition of unlabeled Vpu variants the average distance between residues 39/103 shifted to a much shorter range around 2.5 nm, thereby revealing a major restructuring of Ca^2+^-CaM. In the middle of panel B, reconstructed distance distribution is from the denoised DEER signal. This distance distribution was derived from the original signal (Figure S6).

When soluble Vpu variants, i.e. the FL Vpu and its truncated C-terminal region were added to the CaM solution, substantial fraction of the DEER distance distribution between the spin-labeled residues S39C and A103C in Ca^2+^-CaM has shifted to a short range of distances (Figure 3B, middle and lower panels, Figure S7). This proves that Ca^2+^-CaM has restructured, apparently via formation of a Vpu-Ca^2+^-CaM heterocomplex. The remaining distances, still around 5.3 nm, tell that a fraction of Ca^2+^-CaM remains unbound to Vpu at the protein concentrations used and conditions of our experiment. We estimated the fraction of Vpu-bound Ca^2+^-CaM by integrating the DEER-derived distance distributions and found that slightly more than 50% of distances at 5.3 nm shifted to a shorter 2.5-4.0 nm range upon the complex formation (Figure S8). The shorter distances are consistent with the closed conformation of Ca^2+^-bound CaM in a complex with a protein partner calmodulin dependent kinase fragment^4^ (Figure 3A, lower panel). In this conformation, the long central helix of Ca^2+^-CaM is believed to unravel in its central region, possibly due to its unfolding, to transition to a less structured bent conformation, permitting the two lobes come in contact with about 7 residues of the associated protein^31^ (Figure 3A). Furthermore, the binding of the Vpu variants did not lead to an increase in the DEER modulation depth, which is expected if more than one CaM molecule binds Vpu. This tells that Ca^2+^-CaM binds soluble Vpu monomer rather than its oligomer. The observed moderate decrease of the DEER modulation depth, damped signal oscillations, and resulting broader distance distributions (Figure 3B middle and lower panels, and Figure S9A), are all in line with the increased conformational heterogeneity of Ca^2+^-CaM, a known propensity of the central helix to break in its middle region. It also may indicate that CaM grips and occludes the Vpu region contributing to its oligomerization thereby shifting the equilibrium among Vpu oligomers.

We attempted next to study the conformational changes in Ca^2+^-CaM upon the addition of just the C-terminal region of Vpu (residues 29-81) without the SUMO tag (tag-free Vpu C-terminal region). However, the effect of this Vpu construct on the Ca^2+^-CaM structure was small. We assumed this could be because of strong Vpu C-terminal region homooligomerization, which would hamper its binding to CaM.

### DEER spectroscopy reveals that the monomers in the soluble Vpu oligomer dissociate upon forming a complex with Ca^2+^-CaM

As noted above, the modulation depth of the DEER signals for doubly spin-labeled Ca^2+^-CaM corresponded to the monomeric CaM (just two interacting spins) with and without Vpu present, under the conditions of our experiments (Figure 3). However, we know that at concentrations above 5 µM FL Vpu is oligomeric in solution^19^. Therefore, if the Vpu homooligomer is somehow preserved, it is possible that more than one molecule of spin-labeled Ca^2+^-CaM could be bound to it. In such case, the modulation depth of the DEER signal for the doubly spin-labeled CaM should be much greater than that in the Ca^2+^-CaM monomer, unless the additional distances in such complex are too large to contribute to the modulation depth on the time scale of our DEER experiment. However, as described in the previous section, we did not observe an increase in the DEER modulation depth for the Vpu-bound Ca^2+^-CaM (Figure 3), suggesting that the collective CaM binding is rather unlikely. Nevertheless, we conducted additional experiments to determine the effect of Ca^2+^-CaM on the soluble Vpu homooligomer. We tested this using the FL Vpu having an MTSL attached to the residue L42C in the helix 2. A clear but not oscillating DEER signal in soluble Vpu oligomer was recorded when no Ca^2+^-CaM was present in the sample (Figure 4A and Figure S9B).

**Figure 4.**
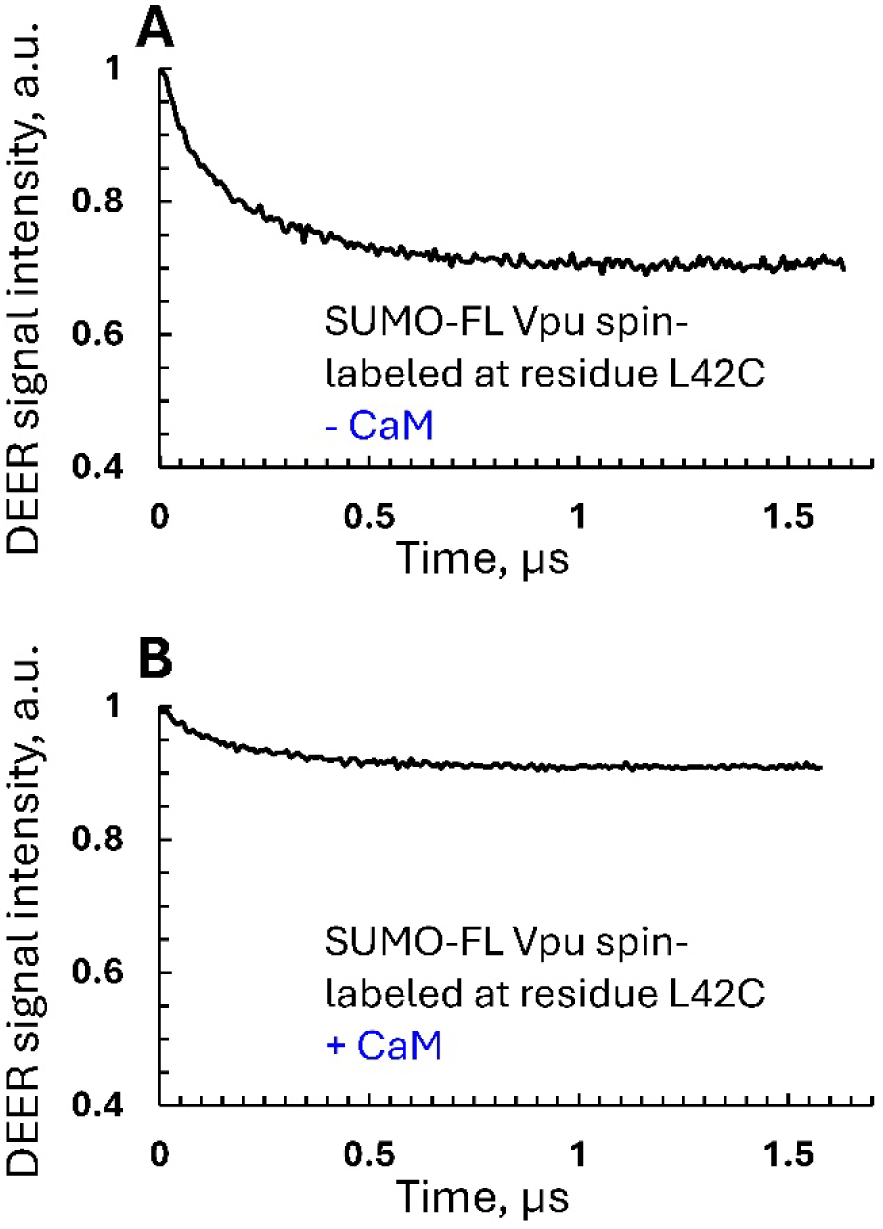
DEER data for soluble FL Vpu (within the SUMO-Vpu construct) spin-labeled at residue L42C. **(A)** Baseline slope corrected DEER signals for soluble Vpu in the absence and (**B**) in the presence of Ca^2+^-CaM, taken in Vpu-to-CaM equimolar ratio. 1 mM CaCl_2_ was added in the buffer for the samples used in (A) and (B). A large reduction in the DEER modulation depth was apparent upon the addition of Ca^2+^-CaM.

Albeit the recorded DEER modulation depth of about 0.3 (Figure 4A) was significantly less than what would be expected for a Vpu hexamer or heptamer^20^ due to the persistent incomplete spin labeling of Vpu and further loss of spin label in the following procedures, the profile of the DEER signal for the spin-labeled at L42C FL Vpu clearly shows a complex broad distance distribution that would be reasonable for a homooligomer with a plurality of spin sites, some of which could be separated by the distances long enough to contribute little to the depth. Interestingly, when we conducted the DEER measurement on a sample containing the same concentration of the spin-labeled at L42C Vpu, but added an excess of Ca^2+^-CaM, we observed significant reduction of the DEER modulation depth (more than twofold) (Figure 4B). This strongly supports the dissociation of the FL Vpu homooligomer. This result indicates that the FL Vpu and Ca^2+^-CaM most likely interact in 1:1 stoichiometry; therefore, the Vpu homooligomer partly disassembles allowing for the formation of the equimolar Vpu-Ca^2+^-CaM heterooligomer.

### DEER spectroscopy shows that the C-terminal region of Vpu undergoes restructuring upon binding to Ca^2+^-CaM

To better understand the Vpu-Ca^2+^-CaM binding mechanism, we studied the spin-labeled double cysteine mutant Q36C/I61C of the Vpu C-terminal region (within the SUMO Vpu C-terminus construct) with cysteines located in the beginning of the helix 2 and in the helix 3 (Figure 5). We measured by DEER the distances between these residues in samples of Vpu C-terminal region with and without Ca^2+^-CaM present (Figure 5A). Remarkably, upon the interaction of Vpu C-terminus with Ca^2+^-CaM, we observed a major shift in the distance between residues Q36C/I61C from about 2.5 nm (no CaM present) to a longer more broadly distributed distances spanning the range of 3.5-5 nm (Figure 5A, upper and lower panels, respectively). This is direct evidence of the Vpu C-terminal region undergoing major conformational transition to conform to the binding site of Ca^2+^-CaM (Figure 5B).

**Figure 5.**
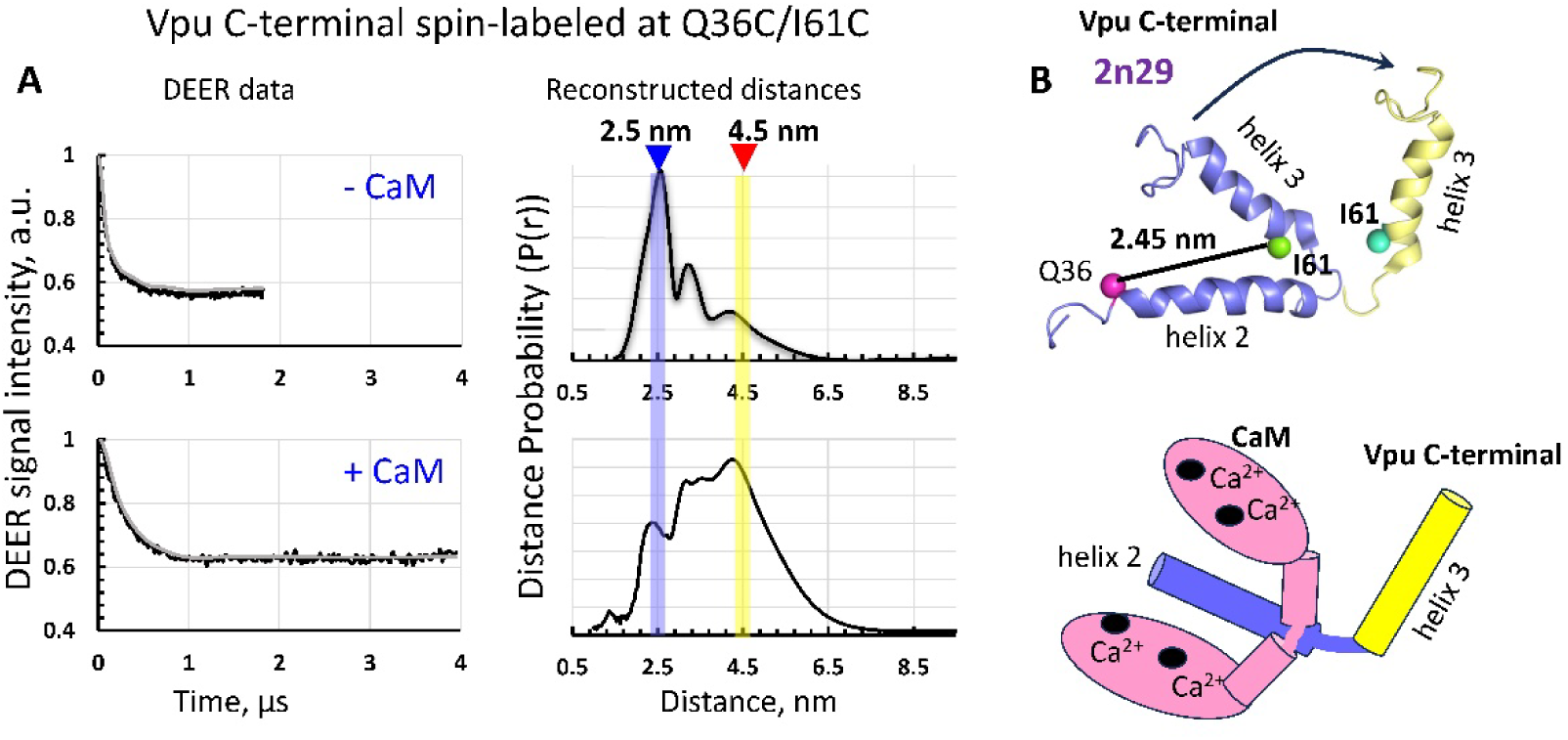
DEER of the spin-labeled at residues Q36C/I61C Vpu C-terminal region confirms significant restructuring upon binding to Ca^2+^-CaM: (**A**) The baseline slope corrected DEER data (left) and reconstructed distances (right) for Vpu C-terminus (with SUMO tag), spin-labeled at residues Q36C/I61C in the absence (upper panels) and the presence (lower panels) of Ca^2+^-CaM. A significant shift in the main distances from about 2.5 nm to 3.5-5 nm was observed, suggesting large restructuring in the Vpu C-terminal region upon binding to Ca^2+^-CaM. (**B**) The proposed conformation rearrangements taking place in Vpu C-terminal region upon binding of Ca^2+^-CaM— the helix 3 moves further away from the helix 2. The ribbon-representation structure (in blue) in the upper panel is that of Vpu C-terminus, as reported by NMR (PDB # 2N29).

It is worth mentioning that the DEER distance between residues Q36C/I61C in the Vpu C-terminal region are very close to the distance of about 2.45 nm predicted based on the solution NMR structure of this protein region^18^ (Figure 5B), thus confirming the arrangement of the helices 2 and 3 in a hairpin, which transitions to a more open conformation upon binding of Ca^2+^-CaM.

The careful examination of the DEER signals (Figure 5A and Figure S9C) shows that the DEER modulation depth (0.4) for the sample of Vpu C-terminus without Ca^2+^-CaM is slightly greater than what is expected for a pair of interacting spin-labels. The value of 0.36 was calculated for the DEER settings of this work, being verified many times with standard biradicals and efficiently labeled globular proteins. This is most likely a result of partial Vpu C-terminus dimerization (Figure S10). However, these Vpu C-terminal oligomers apparently easily dissociate upon the binding to Ca^2+^-CaM because the DEER modulation depth decreased to the expected value (Figure 5A, lower panel, left). This is an additional validation that Vpu and Ca^2+^-CaM bind in equimolar ratio.

Further evidence of direct interaction of the Vpu C-terminal region and Ca^2+^-CaM was obtained by measuring the DEER distances between the singly spin-labeled at residue R37C Vpu C-terminus and the singly spin-labeled at S39C Ca^2+^-CaM (Figure S11).

### FRET spectroscopy confirms that the Vpu C-terminal region has the propensity to self-oligomerize, and this oligomer dissociates upon Vpu-Ca^2+^-CaM complex formation

To further elaborate our studies on the Vpu-Ca^2+^-CaM interactions, we conducted FRET experiments on the tag-free Vpu C-terminus (encompassing residues 29-78), fluorescent-labeled with the Cy3 donor at residue L42C (numbering in FL Vpu), and Ca^2+^-CaM labeled with the Cy5 acceptor at residue S39C. We tested the interaction of this Vpu construct with Ca^2+^-CaM at a low protein concentration below 1 µM, aiming to understand whether in this concentration range the homooligomerization of the Vpu C-terminal region is reduced, compared to >25 µM protein used in the DEER experiments. That is, we expected that the Vpu-Ca^2+^-CaM association could be more efficient at low concentrations. Consequently, we mixed 0.6 µM of the Cy3-Vpu C-terminal region and 0.85 µM of the Cy5-Ca^2+^-CaM to measure the bulk Fӧrster fluorescence resonance energy transfer (FRET) between the Cy3 and Cy5 dyes. In the case where the tag-free Vpu C-terminal and Ca^2+^-CaM form a complex, the Cy5 emission at about 670 nm would be observed due to FRET^32^ (Figure 6A). Indeed, our experiment produced positive results because for the sample of Cy3-Vpu C-terminal region the addition of Cy5-Ca^2+^-CaM produced Cy5 emission at about 665 nm (Figure 6B, red). Such emission band was not observed for the sample of Cy3-Vpu C-terminus in the absence of Ca^2+^-CaM (Figure 6B, black). However, we also noticed that the FRET signal decreased becoming more complex than a simple superposition of just the spectra of unbound and bound species^33^.

**Figure 6.**
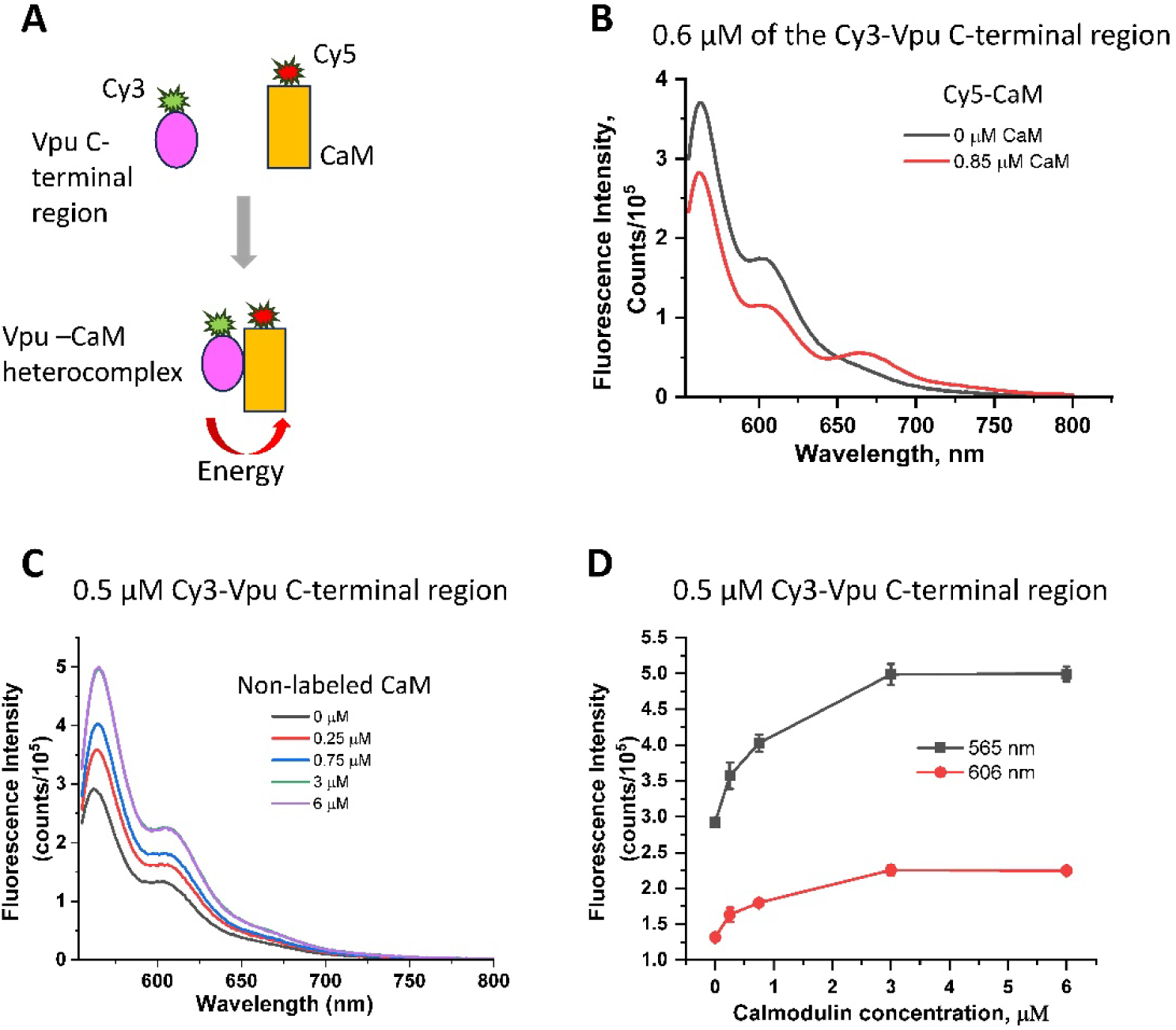
FRET study on the association between the tag-free Vpu C-terminal region and Ca^2+^-CaM: (**A**) Upon mixing the Cy3-Vpu C-terminus and Cy5-Ca^2+^-CaM they form a complex and due to the close proximity FRET occurs between the Cy3 and Cy5 fluorophores. (**B**) The fluorescence spectra of Cy3-labeled Vpu C-terminal and Cy5-labeled Ca^2+^-CaM. Upon the interaction between Cy3-Vpu C-terminal region and Cy5-Ca^2+^-CaM, the emission intensity of the donor at 562nm decreased while the intensity at 665nm of the Cy5 acceptor increases due to FRET. (**C**) The spectra of fluorescently-labeled with Cy3 Vpu C-terminal region under conditions of increasing Ca^2+^-CaM concentration—The intensity of Cy3 fluorescence spectrum increases with the increase of CaM concentration. (**D**) The Cy3-Vpu C-terminus fluorescence intensities at the maximal peak (565 nm) and the shoulder (606 nm) in the presence of different concentrations of Ca^2+^-CaM—Initial fluorescence enhancement was observed and the fluorescence intensity reaches a plateau at 3 µM Ca^2+^-CaM. These data are average of 3 measurements, and the standard deviations are shown.

To explain this, we suggested that there was a third kind of species, most likely homooligomers of the Vpu C-terminal region reported by DEER for a higher protein concentration.

To test this idea, we mixed 0.5 µM of Cy3-labled tag-free Vpu C-terminal region with increasing concentration of non-labeled Ca^2+^-CaM in the range of 0 µM to 6 µM, and recorded the fluorescence spectra of these samples (Figure 6C). We expected that upon increasing the concentration of Ca^2+^-CaM at 0.25 µM, 0.75 µM, 3 µM and 6 µM, a measurable change, most likely increase, in the Cy3 fluorescence would occur due to Cy3-Vpu C-terminal homooligomers dissociation and Cy3-Vpu C-terminal-Ca^2+^-CaM association. It is known that Cy3-biomolecule fluorescence depends on homooligomerization—most often a reduction in the fluorescence intensity is observed due to self-quenching—and binding to a second protein causes fluorescence enhancement^34^. Indeed, the obtained results confirmed our expectations, and we observed Cy3 fluorescence enhancement with fluorescence intensity reaching its maximum in the sample contaning 3 µM non-labeled Ca^2+^-CaM (Figure 6D), presumably because all Vpu C-terminal oligomers were dissociated and monomers were bound to CaM. This result was confirmed by the data from FRET measurements on 0.5 µM/0.5 µM mixture of Cy3/Cy5-labled tag-free Vpu C-terminal region under conditions of increased concentration of non-labeled Ca^2+^-CaM. In this case, we observed reduction in the emission of the Cy5-Vpu C-terminus (Figure S12) at the maximum of its intensity at 662 nm. This points to the dissociation of the Vpu C-terminus homooligomer. These results indicate that the homooligomers were present even at concentrations of the Vpu C-terminal region as low as 0.5 µM. Further studies will provide more details about these homooligomers stability and how other than Vpu concentration factors, such as the presence of solubilization tag, affects this stability.

## DISCUSSION

Despite the CaM-binding motif in the HIV-1 Vpu protein was predicted more than 10 years ago^12^, no experimental evidence of such interaction existed. Moreover, the putative CaM-binding motif contains a substantial region coinciding with the Vpu TM helix 1 (Figure 2). Therefore, it is not immediately clear how the highly soluble CaM interacts with an aa sequence region, which is embedded in the membrane in the case of membrane bound Vpu. The Ca^2+^-CaM regulates several transmembrane proteins^4^ which, however, have CaM-binding sites in their soluble regions like in connexin 45^35^. Our lab has described recently a soluble form of Vpu^19, 20^, which had not been characterized in terms of protein-protein interaction having possible physiological relevance. Therefore, in this work, we sought to test if soluble Vpu can bind Ca^2+^-CaM. Our aa sequence analysis revealed a more extended CaM-binding region in Vpu than the one predicted earlier^12^. This region encompasses residues in the Vpu TM helix 1, helix 2, and the loop between the helices 1 and 2, presenting an ensemble of more than one canonical CaM-binding motif, which is 1-8-14, 1-5-10, and IQ-like motifs, for example (Figure 2). The absence of the unique canonical CaM-binding motif in Vpu is not a surprise, being a common case for other HIV-1 CaM-binding proteins^8, 12^. What is more, it may well be that the Vpu has more than one CaM-binding site and, hypothetically, they are advantageous for the Vpu-CaM interactions. However, to determine this with high certainty, a more detailed study would be necessary. In this line of thought, hybrid CaM-bunding motifs have been identified in cellular proteins as well, and in some cases these sites have cooperative effects^21–23^. Also, it was found that the matrix proteins (MA) from different HIV-1 subtypes differentiate in their interaction with CaM, thus having implications for their biological function^11^. All in all, there is growing evidence that hybrid and non-canonical CaM-binding aa motifs increase the diversity and versatility of CaM-protein partner interactions. Therefore, it may well be that the availability of hybrid binding sites is advantageous for the Vpu function.

With high certainty, however, this work provides robust experimental proof to the fact that the recently identified soluble Vpu forms readily interact with Ca^2+^-CaM. This is based on our comprehensive DEER spectroscopy results on the interaction of spin-labeled Ca^2+^-CaM and Vpu. The results clearly showed that both Ca^2+^-CaM and Vpu undergo substantial conformational changes to conform to the specifics of the CaM-Vpu complex. We concluded that upon binding to Vpu, the Ca^2+^-CaM adopts closed conformation, based on the dramatic change of the DEER-derived distances between spin-labeled residues S39C and A103C, which decreased from 5.3 nm down to 2.5 nm (Figure 3). This major restructuring is in line with previously observed conformational changes when Ca^2+^-CaM interacts with the native for the cell proteins, where in the process the N- and C-terminal lobes of CaM come close to each other wrapping the binding region of the protein partner; the process facilitated by partial unwinding of the central helix^4, 12^. In addition, we found that the Vpu C-terminal region undergoes substantial restructuring to facilitate the interaction with CaM by moving the helix 3 further away from the helix 2 (Figure 5) as the helix bundle would not conform to CaM binding cleft formed by the fraying of the central helix and the destabilized terminal lobes. This Vpu restructuring may be necessary because a more extended helix in the Vpu C-terminal region is needed for the stabilization of the Vpu-Ca^2+^-CaM complex, similar to the transition from disordered to ordered state and helix expansion observed for the SK2 channels^36^. Another plausible reason is that the helices 2 and 3 in the truncated Vpu C-terminal region form a U-shape structure due to electrostatic interaction between the highly positive helix 2 and the acidic helix 3^18^. However, apparently, the binding of Ca^2+^-CaM to helix 2 of Vpu is strong enough to overcome the Vpu helix2-helix3 electrostatic interaction. Indeed, the helices 2 and 3 of Vpu participate in multiple interactions with host proteins^14, 37^. Also, the helix 2 was found to associate with the membrane surface^18^. Therefore, the splicing out of these Vpu helices is not unexpected. Furthermore, one may hypothesize that under physiological conditions the hairpin arrangement of the soluble helices 2 and 3 of Vpu is just a transient state, which is merely populated in the presence of a binding partner. This may warrant a further study.

The association between the Vpu C-terminal region and Ca^2+^-CaM was further supported by the FRET results (Figure 6). Strikingly, we were able to undoubtedly observe FRET between the tag-free Vpu C-terminal region and Ca^2+^-CaM, thus overcoming the challenges for DEER spectroscopy to detect this interaction using higher protein concentrations (>25 µM). We believe this is because of the reduced propensity of the tag-free Vpu C-terminal to form homooligomers at concertation as low as 1 µM or less (Figure 6C and D). Such homooligomerization was observed for the chimera construct of the SUMO-Vpu C-terminal region (Supporting figure S9). However, for the latter construct, DEER spectroscopy was successful in detecting the interaction of the SUMO-Vpu C-terminal region with Ca^2+^-CaM (Figure 3B). This is likely a result of increased solubility of the SUMO-Vpu C-terminal construct leading to less efficient self-association.

Based on the aa sequence analysis and comparison with other CaM-binding protein motifs^3, 4^, as well the DEER and FRET results from our experiments on the spin- and fluorescently-labeled Ca^2+^-CaM and FL or truncated Vpu (Figures 3, 5, and 6), we propose that the helix 2 of Vpu and its flanking regions make a main contribution to the Vpu-Ca^2+^-CaM interaction, aided by the residues in the TM helix 1.

The results from this study also suggest that Vpu and Ca^2+^-CaM interact in equimolar ratio, which is supported by the reduction in the modulation dept of the DEER signal for soluble Vpu homooligomers with and without Ca^2+^-CaM (Figure 4) and the dissociation of the oligomers of the Vpu C-terminal region in the presence of Ca^2+^-CaM detected by FRET (Figures 6 and S).

Here, we propose a model of the interaction of the soluble Vpu with Ca^2+^-CaM (Figure 7), in which the Ca^2+^-CaM “extracts” a Vpu monomer from its homooligomer to form a heterocomplex in a 1:1 molar ratio. We can further speculate that the Vpu-CaM complex may have a role in trafficking the Vpu protein to the membrane site, as proposed for the HIV-1 Gag protein^9^. Future studies will be necessary to clarify this supposition. Vpu is thought to be synthesized at the ER membrane, although specific details of this process have not been documented in existing literature. Furthermore, the protein is found in the plasma membranes, and trans-Golgi network (TGN)^13, 38, 39^, but its trafficking to these membranes is currently insufficiently understood. It may well be that the Vpu-Ca^2+^-CaM complex might take part in the process of Vpu distribution in the cells, particularly because the CaM-binding motif(s) in Vpu is highly conserved (Figure 2).

**Figure 7.**
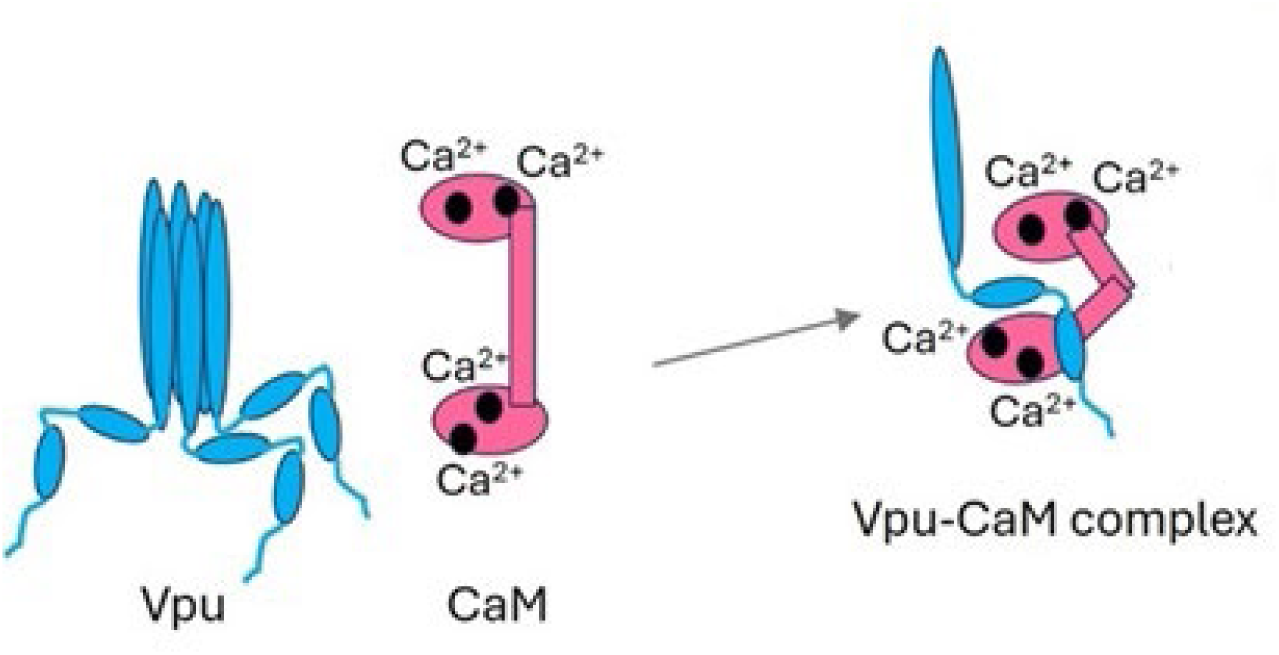
A model of soluble Vpu-CaM complex formation: Soluble Vpu binds to Ca^2+^-CaM in 1:1 stoichiometry; therefore, the soluble Vpu oligomer dissociates. The Ca^2+^ ions bound to CaM are shown as black circles.

This study provides new insight into the interaction of the HIV-1 Vpu protein with cellular components. Studies are being conducted in our lab to understand whether the membrane-bound Vpu also interacts with CaM, or the membrane surface displaces CaM in a competitive interaction with the amphipathic helix 2 of Vpu. In ongoing studies, we also aim to comprehensively characterize the soluble Vpu-Ca^2+^-CaM complex and by detailed DEER mapping to identify with higher certainty the Vpu-CaM binding interface. This insight may aid in designing Vpu inhibitors.

Here, we experimentally detected the Vpu-Ca^2+^-CaM interaction, which adds to the lineage of the previously identified complexes of HIV-1 proteins with CaM. Thus, the growing evidence of the interactions of HIV-1 proteins with CaM might suggest that the virus efficiently utilizes CaM to adjust to the cell environment. Indeed, this might be a rational choice for HIV-1 because of the ubiquitous localization and high concentration of CaM in the cell.

## MATERIALS AND METHODS

### Protein design, cloning, mutagenesis, and expression

All fusion protein constructs, their cloning, and the protein mutations were designed in our lab; the DNA synthesis, cloning, and mutagenesis were conducted by GenScript, Inc.

Full-length (FL) Vpu was designed as a SUMO-Vpu-His_10_ construct (referred here as SUMO-Vpu) having a SUMO tag at the N-terminus and poly-histidine purification tag at the C-terminus (Figure S2). The truncated C-terminal region of Vpu (residues 29-81) was designed as a His_8_-SUMO-Vpu C-terminal construct (referred here as SUMO-Vpu C-terminus or SUMO-Vpu C-terminal region) (Figure S3). The protein solubilization using SUMO and other tags (e.g., MBP) was used in studies of proteins, including membrane proteins and viroporins^40, 41^. Furthermore, SUMO-tagged viroporin (the same family as Vpu) from coronavirus maintained its activity in cells^40^. Therefore, we strongly believe that the SUMO tag does not present a destabilizing factor to viroporin structure. The wild-type (WT) Vpu, which we used, has no native cysteines, therefore the cysteine mutants used in this study were generated on the background of WT Vpu. The SUMO-Vpu had a cysteine residue introduced through the mutation L42C (numbering is that of FL Vpu) which is in the Vpu helix 2, and a double cysteine mutant of the SUMO-Vpu C-terminus Q36C/I61C (numbering in FL Vpu) with cysteines in helices 2 and 3 was generated, and finally a single cysteine mutant R37C in the SUMO-Vpu C-terminus. To verify that these point mutations do not affect Vpu folding, we compared the AlphaFold models of the WT Vpu and the cysteine mutants and insignificant differences in the structures of WT vs. mutant were noticed (Figure S1).

In addition, a peptide encompassing the Vpu residues 29-78 with a cysteine residue at position L42C (numbering in FL Vpu) was commercially synthesized (RS Synthesis).

Two CaM variants used in this study were the single cysteine mutant S39C and the double cysteine mutant S39C/A103C.

All proteins were cloned in pET15b *E. coli* expression vector at the Nco I and BamH I cloning sites, and the resulting plasmids carrying the gene for the protein of interest were transformed into chemically competent *E. coli* BL21(DE3) cells for protein expression. To express the proteins, *E. coli* colonies containing the plasmids carrying the SUMO-Vpu, SUMO-Vpu C-terminus and CaM were grown overnight on LB/agar/100 µg/ml ampicillin (Amp) plates at 37°C. A single colony for each protein was selected to inoculate 200 ml of sterilized LB medium (Lennox) supplemented with 100 µg/ml Amp to produce the cell stock solution. The cells were grown overnight at 37 °C and 180-200 rpm in an incubator-shaker. On the next day, 30 ml of the cell stock solution was transferred into each of larger flasks containing 2 l LB/100 µg/ml Amp and the cells were grown in the incubator shaker at 37 °C and 180 rpm. After 2 h 30 min – 3h, when the OD of the cell solution reached 0.6-0.8, the temperature in the incubator-shaker was reduced to 14-18 °C. After the desired temperature was reached, the protein expression under the control of T7 promoter was induced by adding IPTG (Isopropyl ß-D-1-thiogalactopyranoside) to a final concentration of 0.5-1 mM in each of the flasks. The protein expression was conducted overnight in the shaker-incubator at 14-18 °C and 180 rpm for about 16-20 h.

### Protein purification

All proteins were extracted from the cells using sonication and the soluble fractions were collected. Because all proteins (SUMO-Vpu, SUMO-Vpu C-terminus, and CaM) had His-tags, nickel affinity (Ni-affinity) chromatography was used in all purification protocols. The detailed protocols for the individual purification of each protein are provided below. The compounds used in the purifications were: NaCl (Thermo Scientific), HEPES (4-(2-hydroxyethyl)-1-piperazineethanesulfonic acid) (Sigma), TCEP (*tris*-(2-carboxyethyl)-phosphine) (Gold Biotechnology), chicken egg lysozyme (Roche), PMSF (phenyl-methylsulphonyl fluoride) (Gold Biotechnology), Urea (Sigma-Aldrich), Ni-NTA agarose resin (Qiagen), Imidazole (Sigma-Aldrich), glycerol (Thermo Scientific), Tris base (BioRad), sodium phosphate monobasic (Sigma-Aldrich), sodium phosphate dibasic (Millipore-Sigma), DEAE resin (Cytiva), Phenyl Sepharose resin (Cytiva), CaCl_2_ (Sigma), and β-DDM (*n*-Dodecyl β-D-Maltoside) (Anatrace).

#### SUMO-Vpu purification

We harvested the cells by spinning them down in an Avanti J-15R centrifuge (Beckman coulter; JA-4.750 rotor) at 4,100 RPM (1,880 ×g) for 10 min at 4 °C. We discarded the supernatant and collected and resuspended cell pellets in the resuspension buffer containing 20 mM HEPES pH 7.5 and 200 mM NaCl. Then, we added TCEP, chicken egg lysozyme, and PMSF to the resuspended cell solution to final concentrations of 200 μM, 0.5–0.6 mg/mL, and 1 mM, respectively. We then subjected this solution to sonication using a sonicator (U.S. solid ultrasonic processor) to break the cells open, and we separated the cell debris by centrifugation at 7,000 RPM (5,380 ×g) for 15 min in the same centrifuge (JA-10.100 rotor) at 4 °C. We collected the supernatant containing the soluble fraction of cell lysate and then using ultracentrifugation at 20,000 rpm (27,400 ×g) for 1 h in an Optima XE-90 ultracentrifuge (Beckman Coulter) in 70.1Ti rotor at 4 °C we span down most of the membranes. The supernatant was collected, and urea was added to final 8 M in stirring mode. Then, we added to the protein solution Imidazole (Im) to 20 mM final concentration as a background to prevent the binding of impurities to the Ni-NTA resin.

We utilized Ni-affinity chromatography to purify the SUMO-Vpu protein. For Ni-affinity purification, we incubated Ni-NTA agarose resin (1.5 ml/1 L cell culture) at room temperature (RT) for 1–1.5 h. Afterward, we transferred Ni-NTA agarose resin bound with protein to a gravity column and discarded flow-through containing the unbound material. Then, to elute most of the weakly bound protein impurities, we washed the column first with 12 resin volumes of buffer A containing 50 mM of sodium phosphate buffer, pH 7.4; 500 mM of NaCl, and 5 % (w/v) glycerol supplemented with 70 mM Im, 8 M Urea, 200 µM TCEP. Next, the resin was washed with 3 volumes of buffer A supplemented with 85 mM of Im, 8 M Urea, 200 uM TCEP to the column. Afterward, we eluted the SUMO-Vpu protein using 2.5 resin volumes of buffer A supplemented with 320 mM Im, 8 M Urea, 200 µM TCEP. Next, we concentrated the protein at 3,900 RPM (1,700 ×g) in 10-kDa MWCO centrifugal filters (Amicon®) at 4 °C-10 °C. We gradually refolded the protein by washing out the Urea with buffer A supplemented with 1 mM β-DDM. Prior to binding experiments with CaM, the SUMO-Vpu buffer was exchanged with a β-DDM- and EDTA-free buffer. We measured the protein concentration using nanodrop spectrophotometer (Thermo Scientific) using the calculated extinction coefficient of 13,980 M^-1^cm^-1^ and absorbance at 280 nm. We used this highly pure protein in further experiments.

#### SUMO-Vpu C-terminus purification

The cells were harvested by centrifugation in an Avanti J-15R centrifuge (Beckman coulter; JA-4.750 rotor) at 4,100 RPM (1,880 ×g) for 10 min at 4 °C. The cell pellets were resuspended in buffer A containing 20 mM Tris/HCl pH 7.5, 500 mM NaCl, 5% glycerol. To the cell solution, PMSF, lysozyme, and TCEP to final concentrations of 1 mM, 1 mg/ml and 200 µM, respectively, were added. The unbroken cells and cell debris were removed by centrifugation at 7,000 RPM (5,380 ×g) for 15 min in the same centrifuge (JA-10.100 rotor) at 4 °C. The supernatant containing the SUMO-Vpu C-terminal protein was transferred into 27 ml ultracentrifuge tubes and the membranes plus the heavy impurities were spun down in a Beckman ultracentrifuge Optima XE-90 at 40,000 rpm (110,000 ×g) for 1 hour at 4 °C. The supernatant containing SUMO-Vpu C-terminus was subjected to Ni-affinity chromatography using the Ni-NTA resin. To this end, the preequilibrated with buffer A Ni-NTA resin was mixed with the soluble protein fraction for 1 h at 4 °C under constant agitation. Thereafter, the Ni-NTA resin with bound protein was washed with 10 resin volumes of buffer A supplemented with 50 mM Im to remove weakly-bound protein impurities. The SUMO-Vpu C-terminal protein was eluted with 3 resin volumes of buffer A supplemented with 300 mM imidazole.

To increase protein purity, additional purification step using ion exchange chromatography was added. Since the SUMO-Vpu C-terminus is negatively charged at pH 7.4, we bound the protein to DEAE-Sepharose ion-exchange resin for 1h at 4 °C. Prior to this, the Im-eluted protein was desalted to remove the Im and NaCl completely. Removing NaCl allows protein to bind positively charged DEAE resin. To do so, the buffer was exchanged to 20 mM Tris/HCl pH 7.5, 1 mM EDTA, 5% glycerol (w/v) (buffer B). The protein impurities bound to the DEAE-Sepharose resin were removed using a stepwise NaCl concentration gradient: 2-5 resin volumes of 20 mM and 55 mM NaCl in buffer B were used. Thereafter, the SUMO-Vpu C-terminal protein was eluted using buffer B supplemented with 300 mM NaCl. Protein purity was assessed via SDS-PAGE and WB (Figure S3). Protein concentration was estimated using the calculated extinction coefficient of 8,604 M^-1^ cm^-1^ for the absorbance at 280 nm.

#### CaM purification

We harvested the cells by spinning them down in an Avanti J-15R centrifuge at 4,100 RPM for 12 min at 4 °C. We discarded the supernatant and collected and resuspended the cell pellets in the resuspension buffer A containing 20 mM Tris pH 7.4, 150 mM NaCl, 200 μM TCEP, and 5% glycerol. Then, we added chicken egg lysozyme and PMSF to the resuspended cell solution to the final concentrations of 0.4 mg/mL and 1 mM, respectively. We then broke the cells open using the sonicator. We separated the cell debris and unbroken cells by centrifugation at 7,000 RPM (5,380 ×g) for 15 min in the same centrifuge (JA-10.100 rotor) at 4 °C. We transferred the supernatant to 27 ml ultracentrifuge tubes and span down the membranes in an Optima XE-90 Beckman ultracentrifuge at 40,000 RPM (110,000×g) for 1h at 4 °C. We collected the supernatant containing soluble CaM. We purified the CaM protein using Ni-affinity chromatography: We incubated the protein solution with Ni-NTA agarose resin (1.5 ml/1 L cell culture) under constant agitation at 4 °C for 1h. Afterward, we transferred Ni-NTA agarose resin bound with protein to a gravity column and discarded the flow-through fraction containing the unbound material. Then, to elute most of the weakly bound protein impurities, we washed the column with 10 resin volumes of buffer A containing 20 mM Tris pH 7.4, 150 mM NaCl, 200 μM TCEP, 5% glycerol (w/v), supplemented with 65 mM Im. Subsequently, we added 3 resin volumes of buffer A containing 20 mM Tris pH 7.4, 150 mM NaCl, 200 μM TCEP, 5% glycerol (w/v), supplemented with 320 mM Im to the column to elute the bound CaM. To obtain highly pure CaM, we further subjected the Ni-affinity purified protein to Phenyl Sepharose purification, taking advantage of the exposed hydrophobic residues in Ca^2+^-bound CaM^42^. To this end, we exchanged the buffer of the Ni-affinity purified CaM with a buffer B made of 20 mM Tris, pH 7.4; 150 mM NaCl, 200 μM TCEP, 5% glycerol (w/v), and 5 mM CaCl_2_. We bound this protein to the preequilibrated with buffer B Phenyl Sepharose 6 fast flow (High Sub) resin at 4 °C for 1h under constant agitation. Afterward, we transferred the Phenyl Sepharose 6 fast flow (High Sub) resin with bound protein to a gravity column and discarded the flow-through fraction containing the unbound material. Then we washed the resin with bound CaM with 10 resin volume of buffer B. Next, we eluted the bound CaM with 3-resin volumes of buffer C containing 20 mM Tris, pH 7.4; 150 mM NaCl; 200 μM TCEP; 5% glycerol, and 10 mM EGTA. We collected the eluted CaM and concentrated it at 3,900 RPM in 10 kDa MWCO centrifugal filters at 4 ^°^C. We measured protein concentration with the Nanodrop spectrophotometer using the calculated extinction coefficient of 2,980 M^-1^ cm^−1^. The protein purity was confirmed using SDS-PAGE and Western Blotting (Figure S4). We used this highly pure protein in further experiments.

### Protein spin labeling

The SUMO-Vpu single cysteine mutant L42C (the cysteine is in the Vpu polypeptide, and the numbering is in FL Vpu) and CaM single S39C and double S39C/A103C cysteine mutants were labeled with the MTS spin label (MTSL). MTSL is widely used paramagnetic tag in protein ESR studies^43–45^. The SUMO-Vpu C-terminal single R37C and double Q36C/I61C cysteine mutants were labeled with the 3-(2-Iodoacetamido)-PROXYL spin label (ISL) because when MTSL was added the SUMO-Vpu C-term protein precipitated out, like other viral proteins studied previously^46^. Note that the cysteines are those of the Vpu polypeptide and the numbering corresponds to FL Vpu.

#### SUMO-Vpu C-terminal region labeling

Prior to labeling, the buffer of the purified R37C and Q36C/I61C mutants was exchanged to 20 mM Tris/HCl pH 7.9, 150 mM NaCl, 5% (w/v) glycerol and 50 mM TCEP. Then an aliquot of 50 mM ISL stock solution in acetonitrile was added to the protein solutions for the final protein-to-ISL molar ratio of 1:10. The reaction was allowed to proceed for 3-4 h at 22 °C under constant agitation. Unreacted ASL was removed by washing protein solution with a buffer containing Tris/HCl pH 7.4, 150 mM NaCl, 1 mM CaCl_2_ and 5% (w/v) glycerol.

#### CaM labeling

Prior to spin labeling, the buffer of the purified CaM S39C and S39C/A103 cysteine mutants was exchanged to 20 mM Tris, pH 7.4; 150 mM NaCl; 5% glycerol; and 1 mM CaCl_2_. The CaM concertation was about 100 µM. Then, an aliquot of the 100 mM MTSL stock solution in acetonitrile was added to the CaM mutant solution for the final 1:10 (or 15) CaM:MTSL molar ratio. The reaction was allowed to proceed for 2h 30 min at 22 °C and then overnight at 4°C. The unreacted label was removed by washing the protein solution in a 5 kDa MWCO concentrator with a buffer containing 20 mM Tris/HCl pH 7.4; 150 mM NaCl, 5% glycerol (w/v), and 1 mM CaCl_2_. Finaly, the buffer of the spin-labeled S39C/A103C mutant was exchanged to 20 mM Tris/HCl, 150 mM NaCl, 1 mM CaCl_2_ and about 80% D_2_O instead of H_2_O to increase the MTSL phase relaxation times, allowing us to measure longer distances by DEER spectroscopy (DEER for short)^47, 48^.

### Preparation of samples for DEER spectroscopy

The following samples were prepared for DEER measurements: (i) Ca^2+^-CaM spin-labeled at residues S39C/A103C at 30 µM and 40 µM protein concertation in the buffer containing about 80% D_2_O or 20 % H_2_O. (ii) 30 µM Ca^2+^-CaM spin-labeled at residues S39C/A103C mixed with 40 µM of SUMO-FL Vpu L42C unlabeled mutant. The final sample contained about 50% D_2_O. (iii) 30 µM/25 µ Ca^2+^-CaM spin-labeled at residues S39C/A103C mixed with 35 µM/35 µM of SUMO-Vpu C-terminal R37C mutant non-labeled. The final samples contained about 50% D_2_O. (iv) 30 µM Ca^2+^-CaM spin-labeled at residue S39C. (v) 40 µM Ca^2+^-CaM spin-labeled at residues S39C mixed with 40 µM SUMO-FL Vpu spin-labeled at residue R37C. (vi) 30 µM SUMO-Vpu C-terminal region spin-labeled at residues Q36C/I61C. (vii) 30 µM SUMO-Vpu C-terminal region spin-labeled at residues Q36C/I61C mixed with 50 µM unlabeled Ca^2+^-CaM S39C mutant. (viii) 45 µM SUMO FL Vpu spin-labeled at residue L42C. (ix) 45 µM SUMO FL Vpu spin-labeled at residue L42C mixed with unlabeled 60 µM CaM S39C (in Vpu-to-CaM molar ratio of 0.75).

All samples contained either 20-25% (w/v) glycerol or glycerol-d_8_ as cryoprotectant and all buffers contained 1 mM CaCl_2_.

### DEER spectroscopy experiments and data processing

DEER measurements were carried out at ACERT, Cornell University. The measurements were conducted at 60 K using a home-built Ku-band pulse ESR spectrometer operating in MW frequency band around 17.3 GHz ^49, 50^. The sample solutions were loaded into fused silica 2.0 mm I.D. Vitrotubes® capillaries (VitroCom) and flash frozen for the measurements. A standard 4-pulse DEER experiment^51^ setup was the same as used previously^30^. Briefly, for electron spin-echo detection a π/2-π-π pulse sequence with 32-ns π-pulse width was applied at the low-field edge of the nitroxide spectrum and 16-ns π-pulse at a 70 MHz lower frequency pumped at the central peak. A 32-step phase cycle^52^ was applied for suppressing unwanted coherence pathways contributing to the signal and correcting some instrumental artifacts, leaving only small contributions from the three unremovable dipolar pathways endemic to the 4-pulse DEER sequence. Such contributions caused by dipolar coupling are phase-independent and cannot be removed by phase cycling^53^. Nuclear electron spin-echo envelope modulation (ESEEM) caused by surrounding protons was suppressed by summing up four data traces recorded for each timing point in sequential measurements in which initial pulse separations and the start time of detection were advanced in 9.5 ns steps in subsequent measurements, that is by quarter period of the 26.2-MHz nuclear Zeeman frequency of protons at 615 mT and 17.3 GHz working frequency^54^. For samples that contained deuterated buffers, this usually was unnecessary. The recorded DEER signals were background corrected using log-linear baseline^30^, e.g. as shown in Figure S13. It should be noted that in making the background correction the zero frequency of the Pake doublet was checked to minimize the notch or peak, thereby reducing the baseline uncertainty.

The inter-spin distances between the labels were reconstructed using Tikhonov regularization software^55^ and/or SVD as needed. Most of our data were obtained from samples in low micromolar range, fast relaxing (1-1.5 µs), and not showing oscillatory decays. The reconstructions nevertheless were stable and often similar with both methods, but the second was preferred. In this method some of the original DEER signals were denoised^56^ to facilitate sf-SVD reconstructions^57^ leading to generally more stable distance distributions, *P*(*r*)’s, as a matter of a larger number of singular values. For denoised data in many cases we were able to use all sets of prescribed singular values, but for broadly distributed data we had to stop at just 5-7 values as the reconstruction destabilized fast. The obtained distances are similar when reconstructed from the DEER signal before and after denoising (Figure 3 and Figure S8). Note that in these methods no positivity constraint is used, therefore some reconstructed *P*(*r*)’s may contain small negative contributions that originate from some of unwanted contributions to the data. For example, residual ESEEM contributing to the signal in the opposite phase will be faithfully reconstructed as a narrow negative peak at expected position, when it is not a notch, and it can be ignored or manually removed. Orientation selection, multi-spin effects^29^ and other minor artifacts could also produce negative contributions that may help better recognize their origin, help their removal, and mitigate the reconstruction. The positivity constraint, however, like in MEM^58^ seeded by truncated L-curve Tikhonov distributions may convert these artifacts in some cases into enforced *P*(*r*) distortions; however they are usually rather modest^27^. This method of distance reconstruction used was good enough, as we report distances of up to 5.3 nm (for spin-labeled CaM without Vpu (Fig. 3 B) reconstructed from the low-distortition DEER signals recorded at 4.5 µs dipolar evolution time, which is sufficient for stable distances and distance distributions reconstruction of about 5.5 nm^59^ or even > 6 nm (per DeerAnalysis user manual – 2022). To further validate the DEER distances, we used the DeerAnalysis 2019/2022 freeware, downloadable from Dr. Gunnar Jeschke’s website at ETH Zurich^60^, utilizing the automized option of approximate Pake transformation (APT)^61^. In these analyses, we used the already background corrected DEER data (as depicted in Figure S14 and S15). All distances and distance distributions obtained by DeerAnalysis were expectedly very similar to those reconstructed using the Tikhonov regularization and/or sf-SVD methods (as described above) and are within the reliable range of distances and distance distributions. Thus, the multi-tool approach for DEER distance reconstruction, which we used, confirmed the validity of the DEER distances, which we obtained for the spin-labeled CaM and Vpu under the conditions of this work. As a final remark, the DEER data obtained in this work served rather qualitative purpose compared to the cases of structural triangulation^62–64^, for example.

### Fluorescent labeling of Vpu C-terminal region and CaM

#### Vpu C-terminal rgion

For these experiments, the commercially synthesized peptide of Vpu C-terminal region (residues 29-78 in FL Vpu) (RS Synthesis) with purity of > 95% and received as powder was used. This peptide has a single cysteine residue at position L42C (numbering in FL Vpu). The pure Vpu C-terminal region peptide was dissolved in buffer containing 20 mM Tris pH 7.4, 150 mM NaCl, 5 mM DTT and 5% Glycerol and incubated under constant agitation for about 1h at 22 _°_C. Thereafter the DTT was removed using a NAP-5 desalting column (Cytiva) and buffer was exchanged to 20 mM Tris pH 7.4, 150 mM NaCl, 1 mM CaCl2, 50 µM TCEP, 5% glycerol. The DTT-free peptide, at a final concentration of 60 µM, was aliquoted into two separate 1.5 mL tubes containing 500 µL of peptide solution each. The peptide in one of the tubes was labeled with the donor Cyanine3-maleimide (Cy3) and the other with the acceptor Cyanine5-maleimide (Cy5) fluorophores (Lumiprobe) at a 1:5 peptide-to-dye ratio, and both were incubated for 3 h 30 min at 22 _°_C under constant agitation. Thereafter, the samples were placed at 4 _°_C and incubated overnight under constant agitation. On the next day, the unreacted Cy3 and Cy5 were removed by passing the protein/dyes mixtures 2 times through NAP 5 column (Cytiva). The proteins were in a final buffer of 20 mM Tris pH 7.4, 150mM NaCl, 1 mM CaCl_2_, 50 µM TCEP, and 5% glycerol, which was used in all FRET experiments.

#### Calmodulin

The purified CaM in buffer of 20 mM Tris pH 7.4, 150 mM NaCl, 1 mM CaCl_2_, 50 µM TCEP, and 5% glycerol was labeled with Cy5 at the same 1:5 protein-to-dye ratio, and the remaining procedures were conducted as described in Vpu C-terminal region labeling.

During this incubation, all 1.5 mL tubes contain the proteins and dyes were wrapped in aluminum foil to protect the fluorophores from light exposure, ensuring labeling occurred in a dark environment.

The labeling efficiency was quantified using the NanoDrop One spectrophotometer (Thermo Fisher Scientific), as described^65^. We estimated the Cy3 and Cy5 labeling efficiencies were 21% and 20%, respectively.

### FRET experiments and data analysis

The following samples was prepared for FRET experiment: (i) A sample of tag-free Cy3-labeled Vpu C-terminal region at 0.6 µM (control); (ii) A sample containing a mixture of tag-free Cy3-labeled Vpu C-terminal region at 0.6 µM with Cy5-labeled Ca^2+^-CaM taken at 0.6 µM and 0.85 µM, respectively; (ii) A series of 4 samples containing 0.5 µM/0.5 µM of the tag-free Vpu C-terminal region labeled with Cy3 and Cy5, respectively, which were mixed with increased concentration of unlabeled Ca^2+^-CaM at 0 µM, 0.7 µM, 2 µM and 6 µM. A series of 5 samples containing 0.5 µM tag-free Vpu C-terminal region labeled with Cy3 mixed with increasing concentration of unlabeled Ca^2+^-CaM at 0 µM, 0.25 µM, 0.75 µM, 3 µM and 6 µM. All samples were prepared in triplicates. The samples were incubated at room temperature for 6 minutes to allow protein interaction. The FRET data were collected using FS5 spectrofluorometer (Edinburgh Instruments) with the donor excitation at 550nm (2 nm excitation and emission bandwidth), and emission recorded from 555nm to 800nm. The FRET data were collected for each of the different and the triplicate samples.

For the 0.5 µM/0.5 µM of Cy3-Vpu C-terminus/Cy5-Vpu C-terminus with and without Ca^2+^-CaM, the florescence data for the sample containing 6 µM CaM were taken as a background; This is because under these conditions all Vpu C-terminal region was bound to Ca^2+^-CaM; therefore, no FRET between Cy3-Vpu C-terminus and Cy5-Vpu C-terminus was expected, but the signal contained all the background contributions from bleed-through and acceptor direct excitation. After the background subtraction, the remaining Cy5 fluorescence at the maximum at 662 nm corresponded to the degree of Vpu C-terminal homooligomerization, which was significantly reduced at 2 µM Ca^2+^-CaM (Figure S12).

## Supporting information

Manuscript

## DATA AVAILABILITY

The data obtained through this work will be provided upon reasonable request.

## AUTHOR CONTRIBUTION

OI: experiment, data analysis and interpretation, figures; MMI: experiment, data analysis and interpretation, figures; EH: experiment, figures, data analysis; PPB: experimental design, experiment, data analysis and interpretation, figures, writing the manuscript; AO: experiment, data analysis and interpretation, figures; JCRA: experiment, figures; ERG: conception, experimental design, experiment, data analysis and interpretation, writing the manuscript, figures and artwork, project oversight. All Authors contributed to and approved the final version of the manuscript.

