## Supplementary material for "The soluble state of the HIV-1 Vpu protein forms a complex with Ca^2+^-calmodulin": Manuscript

[#]Equal Contribution

A

MGSDSEVNQEAKPEVKPEVKPETHINLKVSDGSSEIFFKIKKTTPLRRLMEAFAKRQGKEMDSLRFLYDGIRIQADQT
PEDLDMEDNDIIEAHREQIGGSPSGNQGSGLVPRGSSGQGSMQPIPIVAIVALVVAIIIAIVVWSIVIIEYRKILRCR
KIDRLIDRLIERAEDSGNESEGEISALVEMGVEMGHHAPWDVDDLSGHHHHHHHHHH

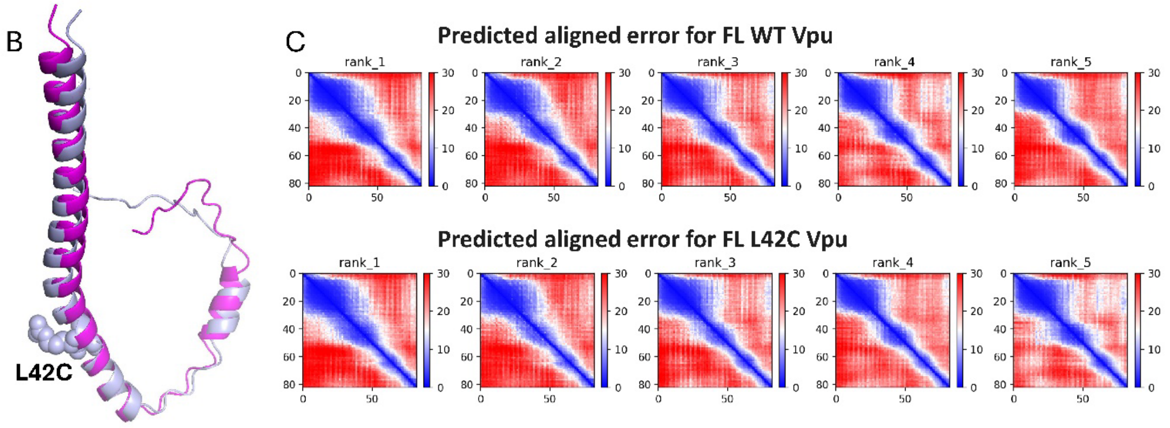


**Figure S1.** Effect of point cysteine mutation L42C on Vpu structure: (A) The amino acid sequence of SUMO-Vpu construct is shown with Vpu's portion highlighted in magenta; (B) AlphaFold-predicted structure of WT and mutant L42C of Vpu –WT protein and the mutant are in magenta and purple-gray, respectively. The atoms of 42C are shown in space-fill representation. There are only slight differences in the predicated structures of WT and L42C mutant of Vpu, which is most likely caused by the dispersion of predicted structures rather than effect of the point mutation. (C) The predicted aligned value for AlphaFold prediction of the structures of WT and mutant Vpu are shown.

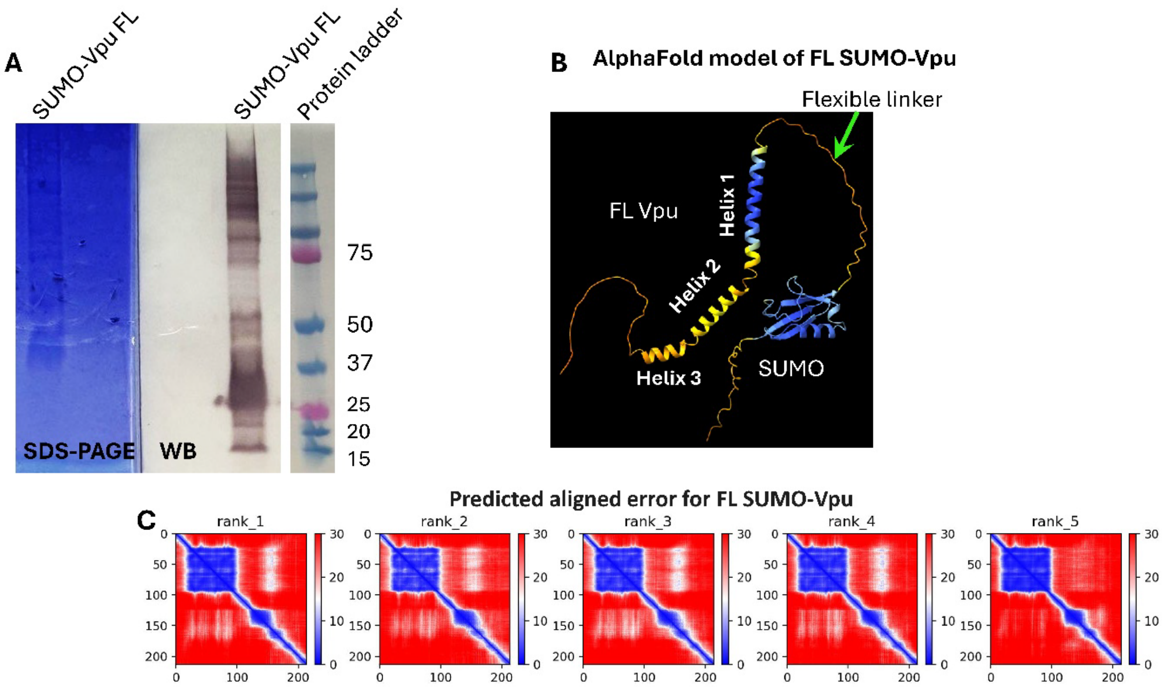


**Figure S2**. The SUMO-Vpu construct used in this study: (**A**) SDS-PAGE and WB of the mutant L42C (numbering in FL Vpu), located in Vpu's Helix 2 in the C-terminal region; (**B**) The AlphaFold model of FL SUMO-Vpu (WT) with the confidence of prediction encoded as blue-to-orange (high-to-low) color range. A long flexible linker connects SUMO with Vpu's N-terminus, which assures that the SUMO tag is far away from the Vpu's oligomerization region (helix 1) and soluble C-terminal region (helices 2 and 3), hence it is unlikely to affect Vpu's homo- and hetero-oligomerization; (**C**) The predicted alignment error for the AlphaFold model in (B).

The SUMO-FL Vpu was purified to a very high purity. The multiple bands visible on the SDS-PAGE and WB are due to oligomerization of the FL Vpu in SDS reported in our past work (Majeed *et al*, 2023, *J Struct Biol*).

**A**

```
MGHHHHHHHHGSNGMSDSEVNQEAKPEVKPEVKPETHINLKVSDGSSEIFFKIKKTTPLR   60
RLMEAFAKRQGKEMDSLRFLYDGIRIQADQTPEDLDMEDNDIIEAHREQIGGSPSGNQGS   120
GLVPRGSSGQGSGMEYRKILRCRKIDRLIDRLIERAEDSGNESEGEISALVEMGVEMGHH   180
APWDVDDL   188
```

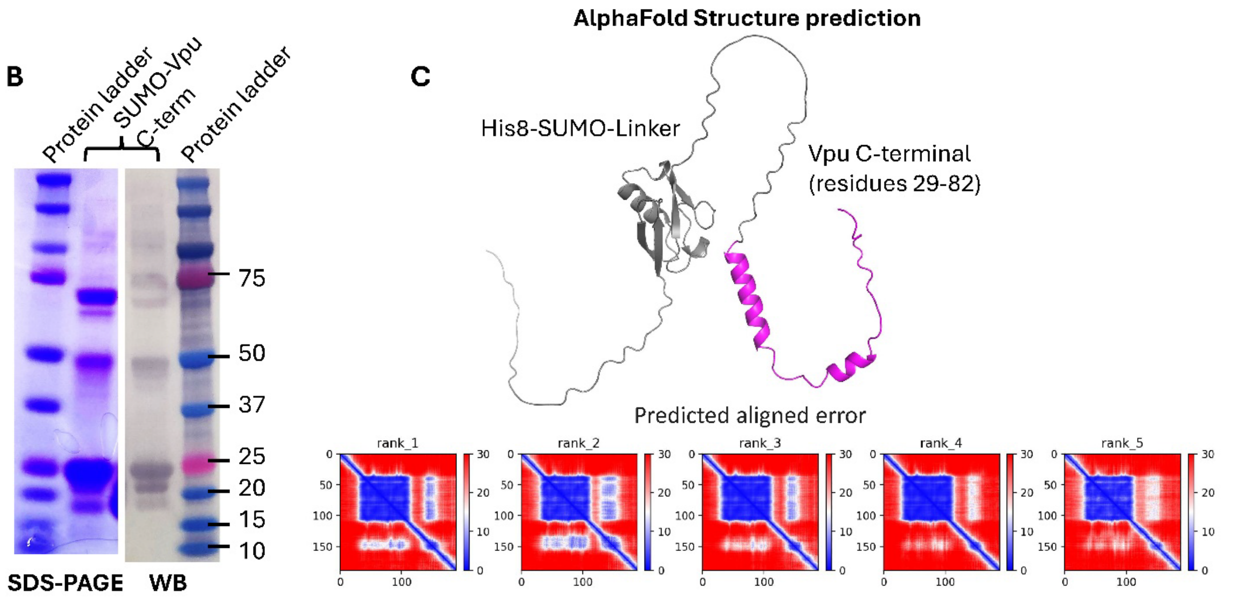


**Figure S3.** SUMO-Vpu C-terminal region construct: (**A**) Amino acid sequence of the FL fusion protein; Vpu's C-terminal region is highlighted in magenta (This sequence carries Q36C mutation). (**B**) SDS-PAGE and western blot (WB) of the purified SUMO-Vpu C-terminal construct—monomeric protein at molecular weight of about 21 kDa and oligomeric protein with higher molecular weight is present. (**C**) AlphaFold predicted structure of the FL SUMO-Vpu C-terminal construct is shown—Vpu's C-terminus is colored in magenta. The linker region between SUMO and Vpu's C-terminus is quite long, making it unlikely to have any significant effect on protein interactions of Vpu's C-terminus.

The SUMO-Vpu C-terminal region was purified to a very high purity. The multiple bands visible on the SDS-PAGE and WB are due to oligomerization of the Vpu C-terminal region, similarly to FL Vpu here (cf. Figure S2) and in our past work (Majeed *et al*, 2023, *J Struct Biol*).

In all cases, the structures of SUMO-Vpu variants were generated using AlphaFold integrated in USF Chimera X Version 1.9 (Yang, Z., et al. "Enhancing UCSF Chimera through web services." *Nucleic Acids Research* 42.W1 (2014): W478-W482), as instructed in the tutorial (https://www.youtube.com/watch?v=le7NatFo8vI).

**A**

```
MGHHHHHHHHHHGLVPRGSGQMADQLTEEQIAEFKEAFSLFDKDGDGTITTKELGTVMR   60
CLGQNPTEAELQDMINEVDADGNGTIDFPEFLTMMARKMKDTDSEEEIREAFRVFDKDGN  120
GYISCAELRHVMTNLGEKLTDEEVDEMIREADIDGDGQVNYEEFVQMMTAK  171
```

**B**

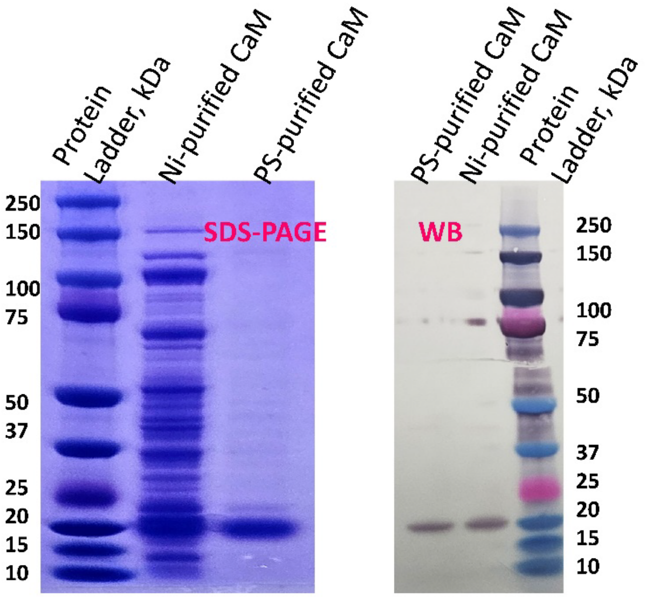


**Figure S4.** Calmodulin (CaM) protein used in this study: (A) The amino acid sequence of the His-tagged CaM construct; (B) SDS-PAGE and western blotting (WB) of CaM at different stages of purification – highly pure protein was obtained after the consecutive Ni-affinity and Phenyl Sepharose (PS) purification. The PS-purified at very high purity CaM was used in this study.

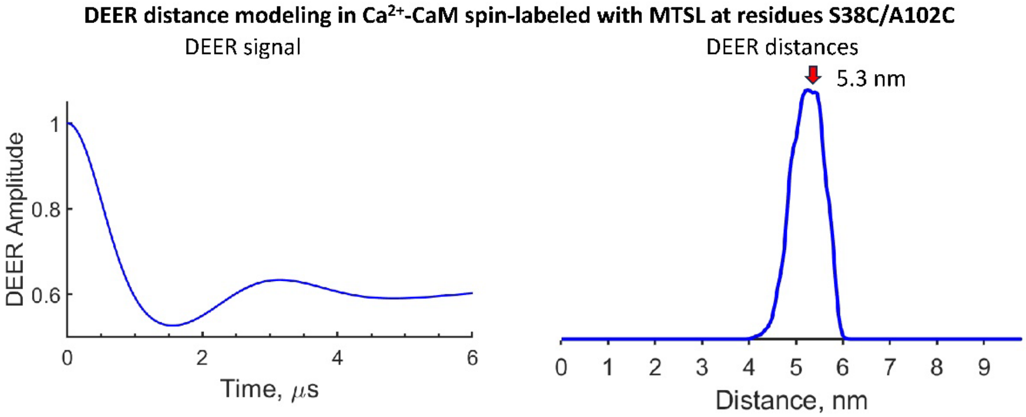


**Figure S5.** Modeled DEER signal and DEER distances between spin-labeled residues S38C/A102C in $Ca^{2+}$-CaM. The average distance of about 5.3 nm was obtained, which is very close to the experimental DEER distance of 5.3 nm shown in Figure 3 in the main text. The molecular modeling software MMM applied to the X-ray structure of CaM (PDB#1exr) was used.

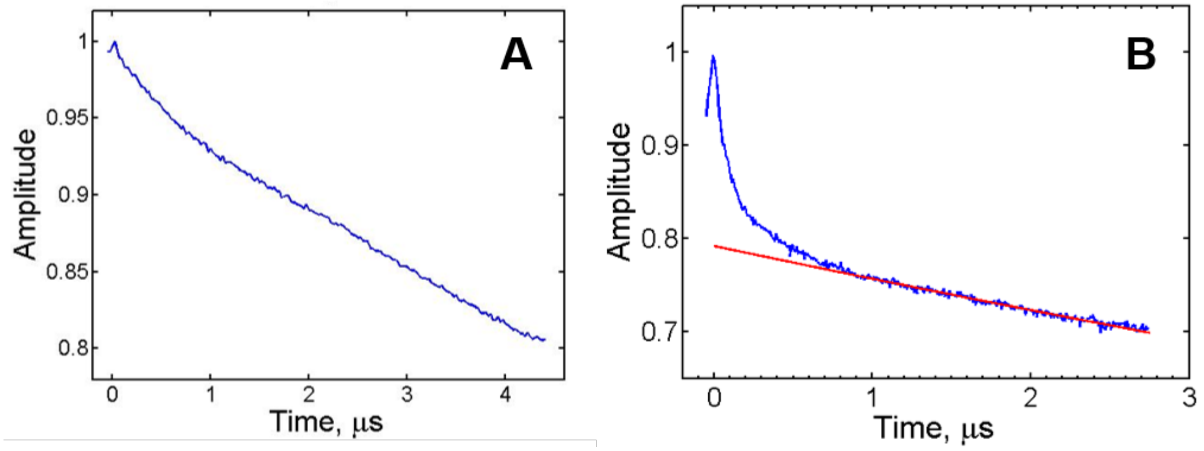


**Figure S6**. The raw DEER data (A) for $Ca^{2+}$- CaM singly spin-labeled at the residue S39C. The concentration of CaM was 32 μM after adding 20% (w/v) glycerol. (B) the DEER data for this construct in the presence of single-spin labeled SUMO - Vpu C-terminus / R37C. The data for the CaM alone shows very shallow DEER modulation pointing to a minor contribution from unknown nature species with long ~5.0 nm distance. The modulation depth is too small (~0.02) for having a noticeable effect on the signal in B. It is unlikely that CaM could form homodimers even in a small fraction. It is conceivable, however, that a minor population of MTSL label could be trapped by binding to the hydrophobic pocket of $Ca^{2+}$ – CaM globular domain.

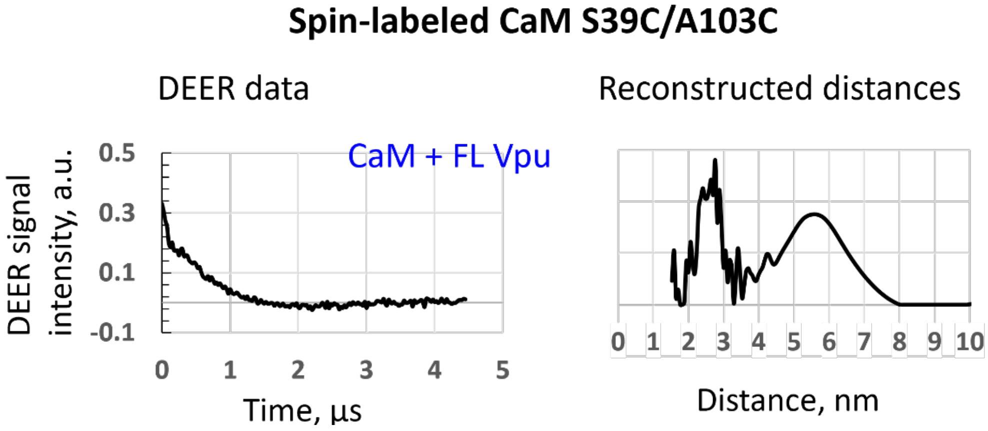


**Figure S7**. **B**aseline-corrected DEER plotted as dipolar modulation (left) and reconstructed distances for the double spin-labeled $Ca^{2+}$-CaM mutant S39C/A103C in the presence of SUMO–FL Vpu. The final concentration after the addition of 20% (w/v), glycerol-$d_8$ to the CaM and Vpu concentrations were 24 µM and 32 µM, respectively.

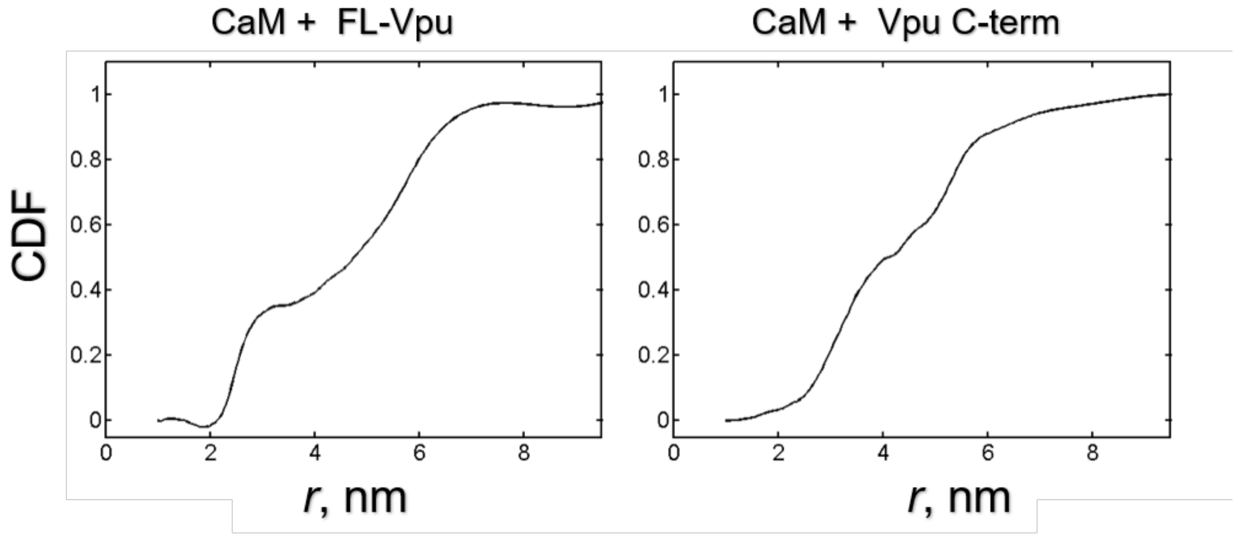


**Figure S8.** The cumulative distribution function (CDF) $\Phi(r)=\int_0^r P(x)dx$ calculated for distance distribution $P(r)$ reconstructed from (left) the DEER data for double spin-labeled $Ca^{2+}$–CaM S39C/A103C in the presence of SUMO-FL Vpu and (right) SUMO-Vpu C-terminus. Greater than 50% of the distances are shorter than 5 nm. The respective $P(r)$'s are used in Figure 3 B.

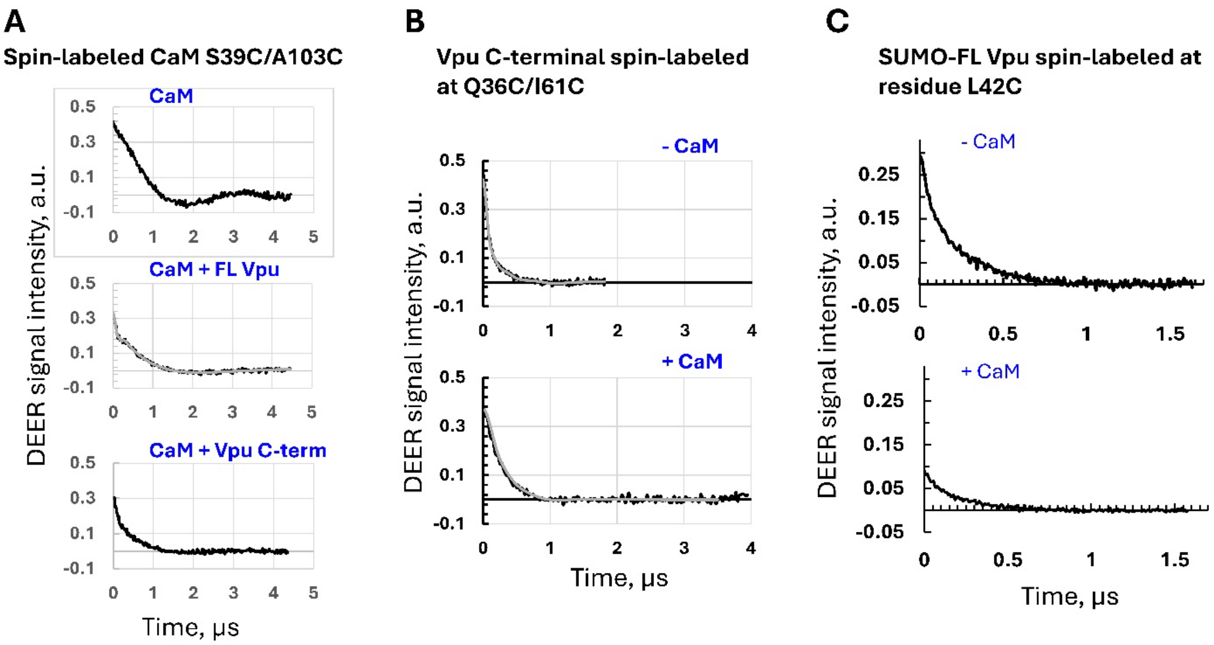


**Figure S9.** Baseline-corrected DEER data plotted as dipolar modulations for: (A) Spin-labeled at residues S39C/A103C $Ca^{2+}$-CaM w/wo Vpu; (B) Spin-labeled at residues Q36C/I61C Vpu C-terminal region w/wo $Ca^{2+}$-CaM; (C) spin-labeled at residue L42C FL Vpu with and without $Ca^{2+}$-CaM. The data are the same as in Figures 3, 4 and 5, but plotted such that the modulation depth can be read at the zero time after shifting to zero and normalization to the scale with the unity being 100% modulation.

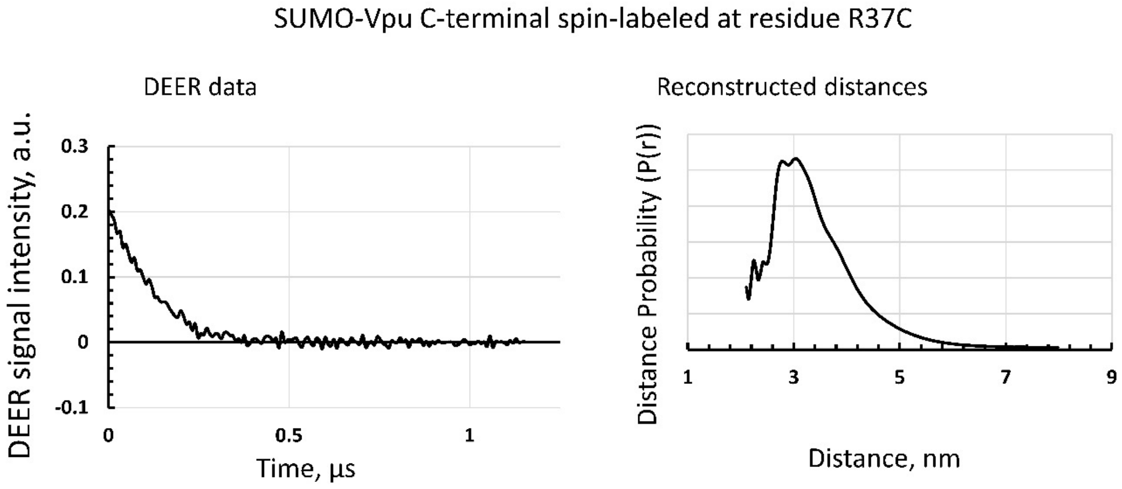


**Figure S10.** DEER data and reconstructed distances for the single spin-labeled at residue R37C (in helix 2 of Vpu) SUMO-Vpu C-terminus. The DEER data indicates that a fraction of the protein shows some form of aggregation, possibly as dimers, yielding a range of distances with a maximum at about 3 nm.

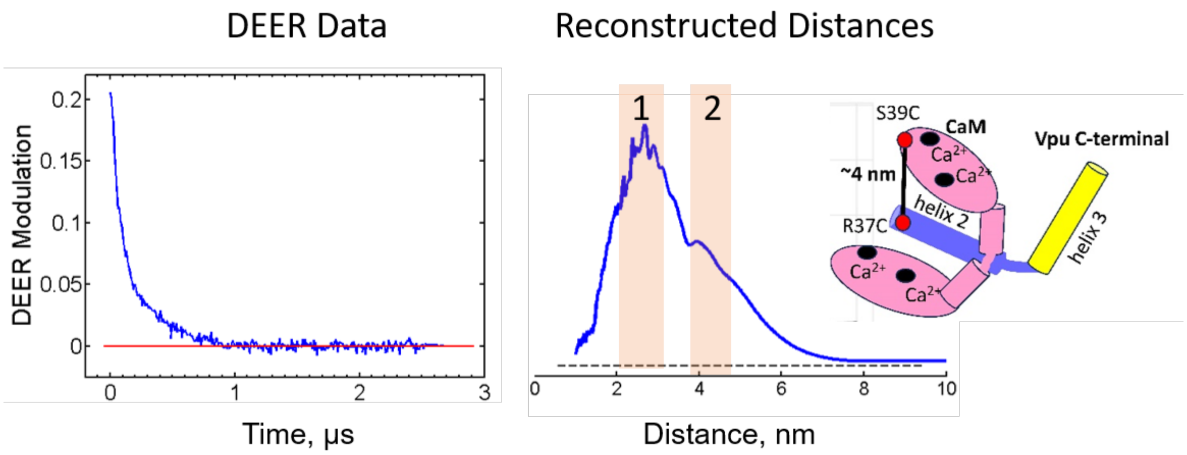


**Figure S11**. (left) Baseline-corrected DEER modulation data for the mixture of spin-labeled Vpu C-terminus at R37C and CaM at S39C. The bimodality is seen by the naked eye. (middle) The reconstructed distances from this data, indicative of a mixture of Vpu C-terminus oligomers (possibly, dimers) with expected distance of about 2.5-3 nm (similarly to the distances found for FL-Vpu in Figure S9) and Vpu C-terminus–CaM heterodimers with not well-resolved distances around 4.2 nm, both present in the sample. The model of Vpu C-terminus–CaM interaction is suggested as one where helix 3 of Vpu moves further away from the helix 2 while $Ca^{2+}$-CaM adopts a more closed conformation to embrace the Vpu helix 2 at the position where the CaM-binding motif of Vpu is located.

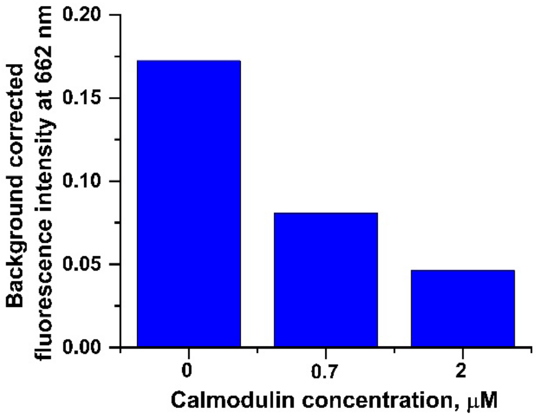


**Figure S12.** Background-corrected fluorescence intensities at 662 nm for the tag-free Vpu C-terminal region labeled with Cy5. The observed Cy5 fluorescence is a result of the FRET between Cy3-Vpu C-terminus/Cy5 Vpu C-terminus in homooligomrs. The fluorescence intensity decreases upon increasing the concentration of $Ca^{2+}$-CaM because of Vpu-CaM heterocomplex formation and Vpu-Vpu homooligomer dissociation. The results were obtained as described in the experimental procedures.

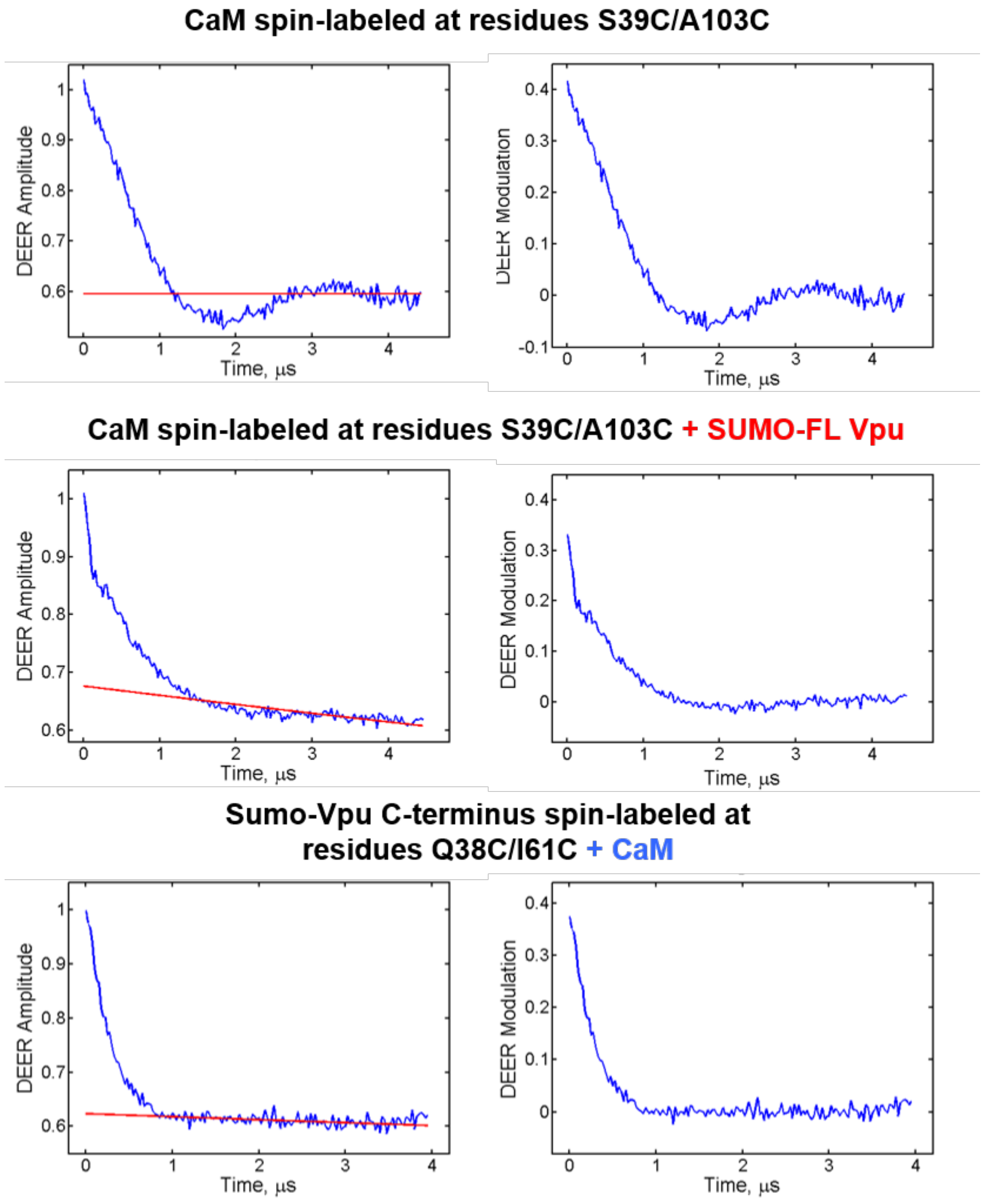


**Figure S13.** Baseline correction examples for the raw DEER data in three representative cases. In the left panels DEER data are plotted together with the log-linear baseline fits (red) except the top panel where constant value had to be used due to insufficient record length. The respective background-subtracted DEER data are shown in the right column. Small "2+1" artifacts are visible is at the record ends.

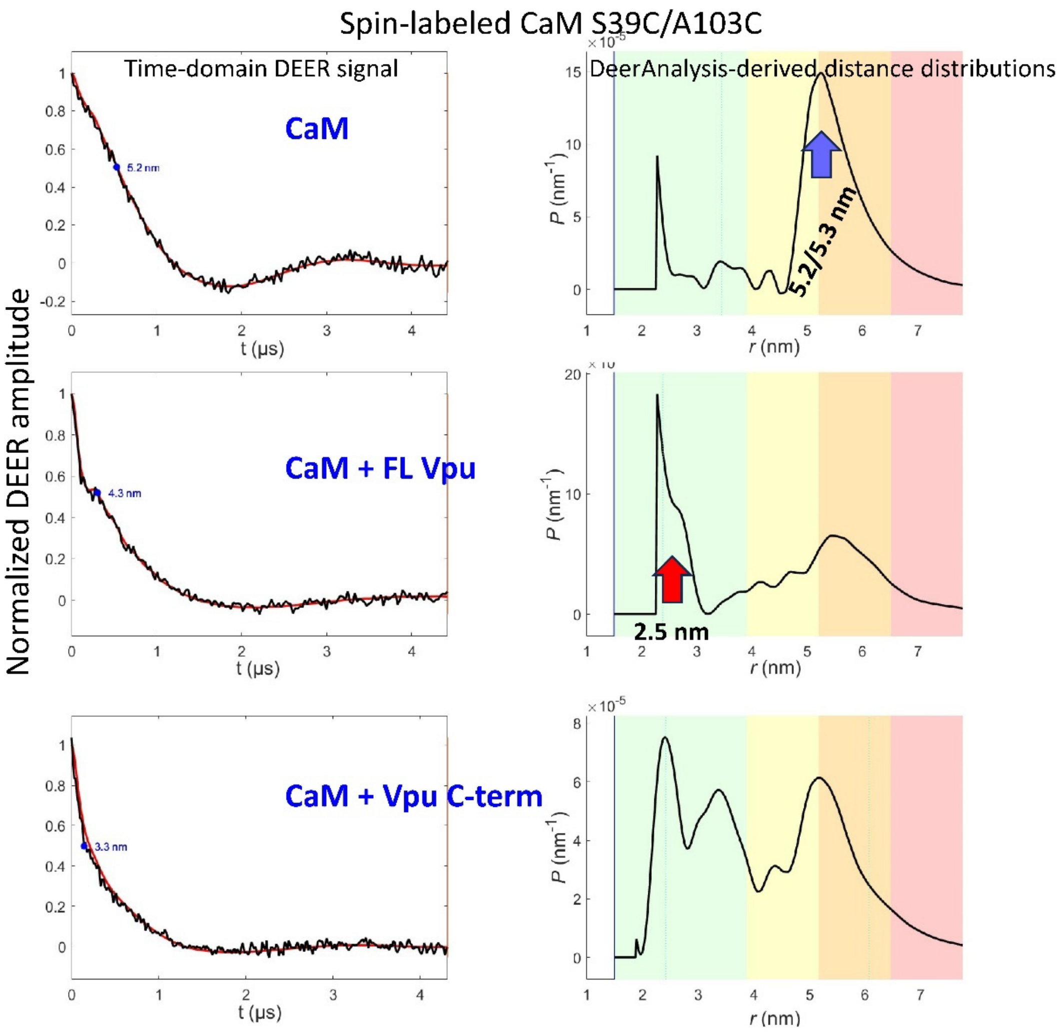


**Figure S14.** DEER distances reconstruction using DeerAnalysis software: Data for spin-labeled $Ca^{2+}$-CaM without Vpu (upper panel), $Ca^{2+}$-CaM in the presence of FL Vpu (SUMO-FL Vpu construct) (middle), and $Ca^{2+}$-CaM in the presence of Vpu C-terminal region (SUMO-Vpu C-terminus construct) (bottom) are shown. The baseline-corrected normalized to unity time-domain DEER data are on the left, and the reconstructed distances using DeerAnalysis 2019 are on the right.

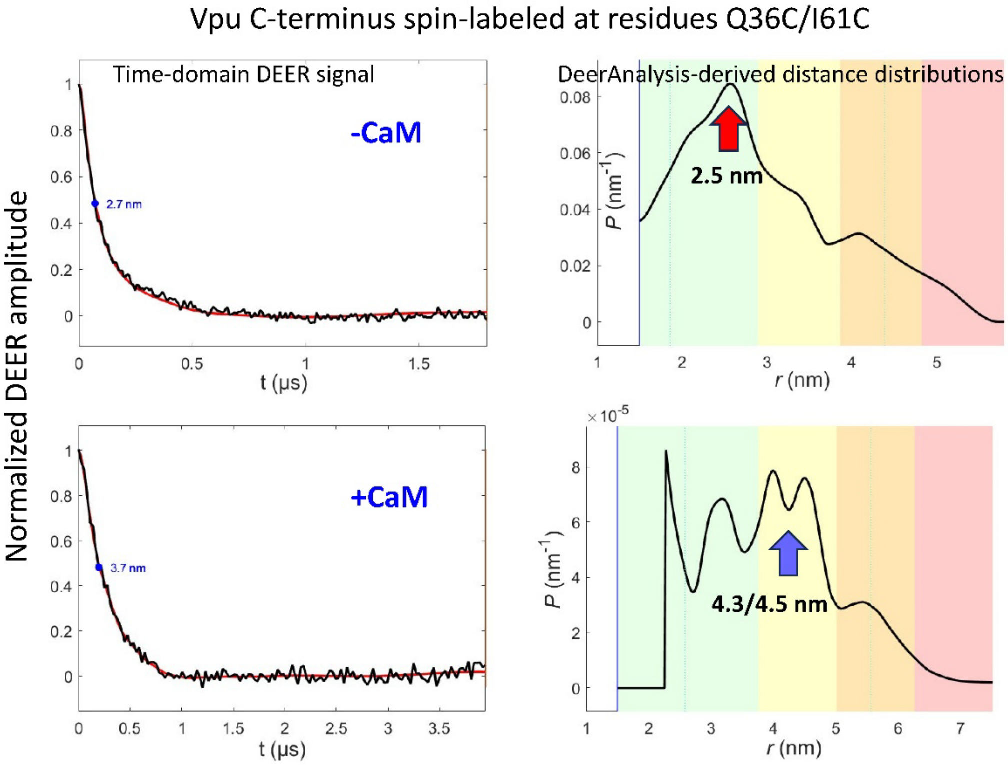


**Figure S15.** DEER distances reconstruction using DeerAnalysis software: Data for spin-labeled Vpu C-terminal region without (upper) and with $Ca^{2+}$-CaM are shown. The baseline-corrected normalized to unity time-domain DEER data are on the left, and the reconstructed distances using DeerAnalysis 2019 are on the right.

In Figures S14 and S15, the color coding for reliability ranges is as follows: Pale green: Shape of distance distribution is reliable. Pale yellow: Mean distance and width are reliable. Pale orange: Mean distance is reliable. Pale red: Long-range distance contributions may be detectable but cannot be quantified. (per DeerAnalysis manual). Our distances are well within the reliable range. Importantly, the distances obtained using DeerAnalysis are very close to those reconstructed using the Tichonov regularization and sf-SVD methods shown in Main Text, Figures 3 and 5.
